# Isolation of thermophilic plastic-degrading bacteria from hot springs of Aotearoa-New Zealand

**DOI:** 10.64898/2026.05.26.727897

**Authors:** Nicholas B. Dragone, Hazel Clemens, Sophie van Hamelsveld, Louise Weaver, Ali Reza Nazmi, Matthew B. Stott

## Abstract

Geothermal springs are unique environments that harbor diverse populations of microorganisms. As a result of their environmental and geochemical variability, different springs can support distinct heterogeneous communities of organisms with unique functional adaptations and metabolic capabilities. A recent molecular survey of Aotearoa-New Zealand hot springs indicated that these springs may support thermophilic microorganisms be able to degrade plastics. To test this, we applied a cultivation-centered approach via an *in situ* enrichment of putative plastic-degraders using high surface are polyethylene terephthalate (PET), polylactic acid (PLA), and polyhydroxybutyrate (PHB) substrates in a diversity of New Zealand hot springs. The plastic associated microbial communities were characterized via marker gene and analyses. Finally, plastic-associated biofilms were used as inoculum to isolate thermophilic plastic degraders. Via this process, we confirm that there are plastic degrading bacteria are present in springs across Aotearoa-New Zealand. Moreover, we isolated two PHB degrading strains (*Cuprividus sp.* and *Rubrobacter sp.)* and demonstrated their capability to metabolize plastic under thermophilic conditions in vitro. While the pathways identified in our plastic degrading isolates suggest they may be able to metabolize plastics for carbon, the primary use of plastics by geothermal microbial communities does not appear to be as an energy source. Instead, they appear to mainly serve as surfaces for microbial attachment, composed primarily of non-plastic degrading taxa.

## Introduction

Plastic pollution remains a major global crisis, with roughly 80% of plastic waste landfilled, incinerated, or entering the natural environment (1), where it persists for decades to centuries and accumulates across terrestrial and aquatic ecosystems, causing long term ecological harm (1–3).Microorganisms and their enzymes have been proposed as a potential tool for breaking down plastics (4), offering a way to improve recycling efficiency and limit microplastic release. Although several hundred putative plastic-degrading microbes have been identified to date (5), none have yet produced an effective, scalable biodegradation method (6). These systems are typically vulnerable to contamination, and most plastic degrading organisms break down polymers far too slowly for industrial use (6). In theory, thermophilic plastic-degrading, microorganisms that remain active at temperatures above 50 °C, should offer significant advantages (7–9). Firstly, plastics become more molecularly mobile at elevated temperatures (above their glass transition temperature (10)), which should enhance enzymatic access, increase degradation rates, and improve substrate breakdown (7, 8, 11). Secondly, high temperature microbial systems should also be more thermodynamically efficient (12, 13), and finally, thermophilic microbial communities should be far less susceptible to contamination by mesophilic organisms (7, 11). Indeed, recent work has suggested that geothermal spring microorganisms may be a valuable source of thermostable enzymes for plastic degradation (14). However, despite the isolation of several thermophilic plastic-degrading microbes (5, 8), we still know remarkably little about how thermophiles interact with plastics in their natural environments. Many fundamental questions remain unanswered, and substantial work is still needed before we can design effective high-temperature microbial plastic-recycling systems based on the activity of thermophilic microorganisms.

Aotearoa-New Zealand (ANZ) is the ideal location to investigate whether microorganisms are capable of investigating thermophilic plastic-degradation. ANZ contains a variety of geothermal zones – most notably the Taupō Volcanic Zone (TVZ) which spans 8000 km^2^ across New Zealand’s North Island and has thousands of geothermal springs formed by oceanic crust melting and dewatering associated with the subduction at the Pacific-Australian tectonic plate boundary (15). As a result of variations in subsurface deposit structure, high heat flux, and deep convection of groundwater, surface features of the TVZ are highly variable and span wide range of physiochemical conditions (16). Extensive biological characterization across these springs has revealed heterogeneous microbial communities, novel diversity, and unique functions (16–19). Moreover, cultivation-independent-based evidence from a limited number of springs from Kuirau Park, Rotorua has suggested that the TVZ’s geothermal springs may contain a large number of putative plastic degrading taxa (14). Based on this evidence, and due to the environmental stress faced by organisms living in these surface features (i.e. nutrient stress, temperature stress), we proposed that we may be able to show that at least some of the organisms found across the heterogeneous microbiomes of these springs are capable of degrading plastic.

To evaluate the potential of Aotearoa-New Zealand’s geothermal springs to harbor thermophilic plastic degraders, and to address how microorganisms interact with plastic introduced into hot springs, we incubated three plastics (polyethylene terephthalate (PET), polylactic acid (PLA), and polyhydroxybutyrate (PHB) in four TVZ springs and one hot spring feature in the South Island of ANZ that spanned a range of physiochemical conditions (pH 5.2 – 8.2, temp 45.0 – 83.8 °C) and geographic distance (<5 km - 550 km). After an up to six month *in situ* incubation, we assessed the sessile microbial communities associated with our plastic substrates via a cultivation-independent marker gene sequencing to evaluate how the introduction of plastic may influence microbial community composition. Using these data we then undertook targeted microbial culturing and whole genome sequencing to evaluate whether any organisms isolated from geothermal springs are capable of degrading plastic at >50 °C *in vitro*. Finally, we used predictive modeling, based on previous extensive characterization of geothermal spring microbial communities and environmental conditions (16), to identify springs with the high potential to contain thermophilic plastic degraders that could be explored in the future.

## Methods

### 3D printed plastic substrates

3D printed substrates were used to as a substrate for organisms to colonize. Triply Periodic Minimal Surface (TPMS) gyroids- a complex mathematically defined shapes that repeats in three dimensions – were designed using TPMS studio ((20); see Figure S1 for the TPMS gyroid design). Three different polymers were used to print the gyroids, polyhydroxyalkanoates, poly lactic acid and polyethylene terephthalate glycol (Figure 2). TPMS gyroids were manufactured using 3D printing with BambooLabs 3D printer using the products: “All PHA Natural” (colorFabb, USA), “PLA+” (eSun industries ltd, China) and “PETG-lite” filaments (eSun industries ltd, China). PETG is an amorphous thermoplastic copolymer resulting from addition of glycol to PET making it more suitable for 3D printing.

**Figure 1:**
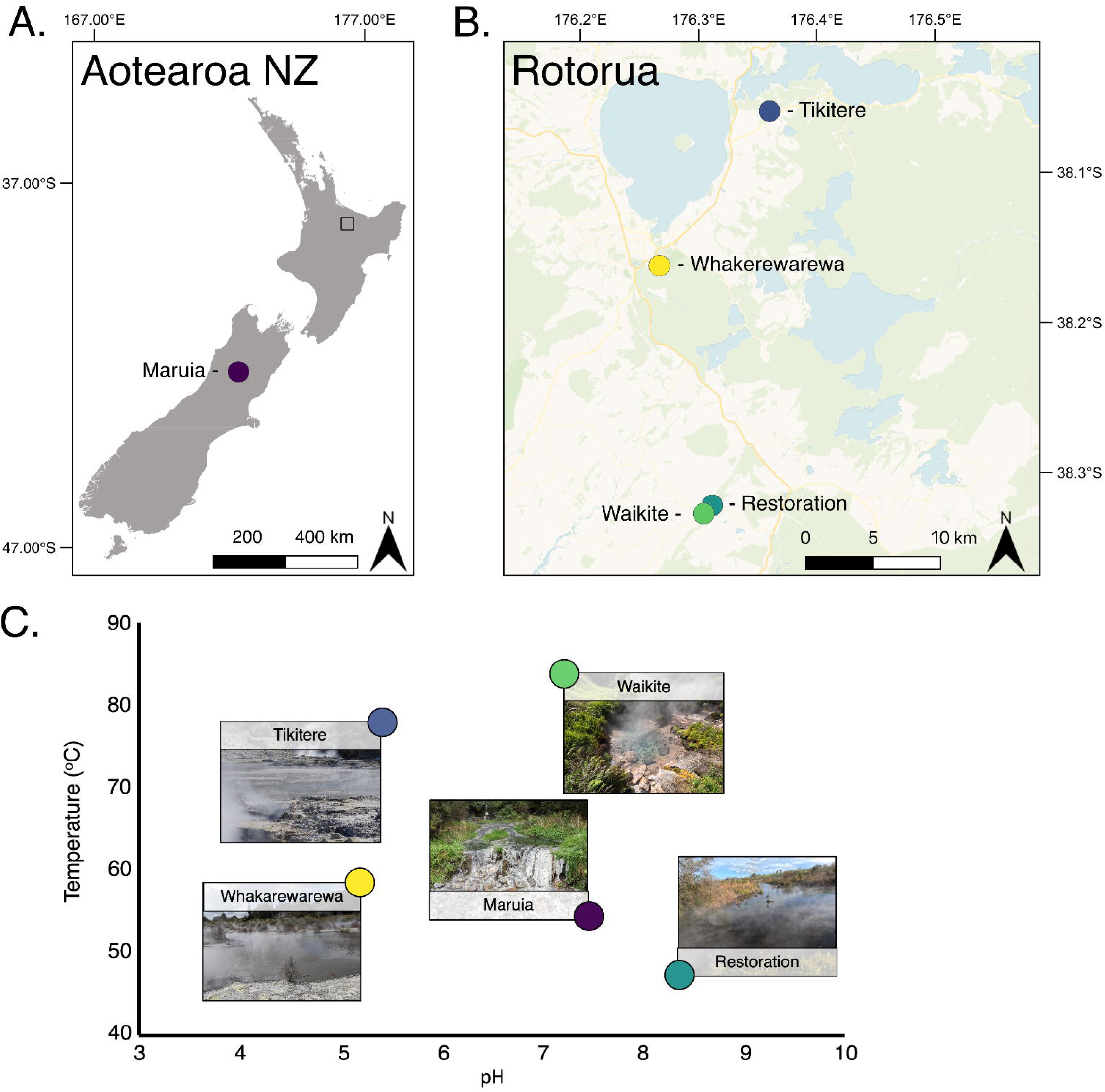
Spring locations. Geothermal spring locations are presented as colored circles across all three images (Maruia; black, Tikitere; blue, Whakarewarewa Village; yellow, Waikite; green, Waikite Restoration area; teal) **A)** Map of Aotearoa New Zealand showing the location of Maruia spring (South Island) and inset indicating the region of the North Island where the other four springs were located. (**B)** Map of Rotorua and surrounding areas detailing indicating the location of the four Taupō Volcanic Zone springs. **C)** Primary conditions of the five springs used for the *in-situ* incubations. Spring temperature and pH reported here are from the start of the multi-polymer incubation, additional measurements can be found in Dataset S1.

**Figure 2:**
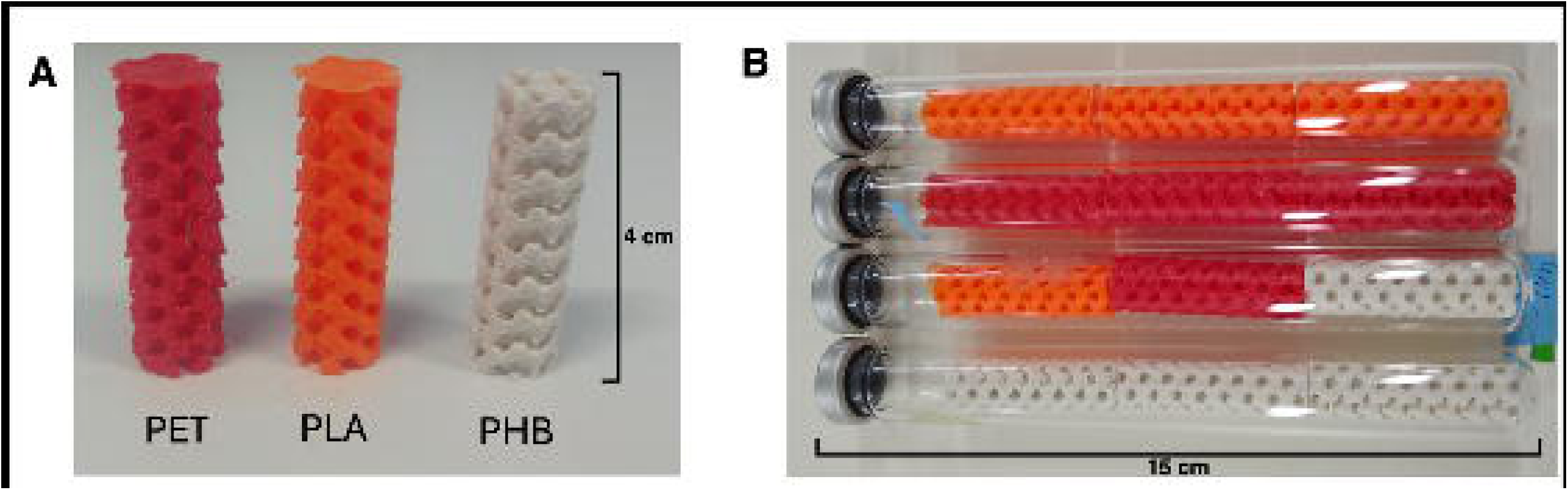
A) Polyethylene terephthalate (PET), Polylactic acid (PLA), and Polyhydroxybutyrate (PHB) gyroids used for the time series and multi-polymer incubations. **B)** Experimental chambers for the multi-polymer incubation. Onsite, holes were cut in the top of the chambers and target hot spring water added to displace the air headspace. The chambers were then suspended in place in via cable ties and heavy gauge nylon fishing line.

The 3d printing stocks contained several additives, though the quantities and composition were challenging to ascertain. According to the producers and material safety data sheets (MSDS): “All PHA Natural” filament are 100% biodegradable, though the details of the mixture of PHAs is not disclosed; the “PETG-lite” filament are 100% PETG with color pigments added; “PLA+” filaments have ∼1.8-2.9% calcium carbonate and 5% of a an additive described in MSDS as: Graft polymer of allylmethacrylate / butylacrylate / decamethylcyclopentasiloxane / 3-[dimethoxy(methyl)silyl]propylmethacrylate / hexamethylcyclotrisiloxane/ methylmethacrylate / octamethylcyclotetrasiloxane.

### Field incubations

Five hot springs across a range of physicochemical conditions and geographic distances were chosen to host our plastic enrichments. Additional information about the incubation experiment and the geochemical properties of each spring can be found in Figure 1 and Dataset S1. Sterilized glass anaerobic culturing tubes with butyl stoppers (Belco Glass, NJ, USA) were used as incubation chambers for the 3D printed substrates (Figure 2, Figure S2). Experimental tubes (with contents exposed to spring water) had holes cut through the seal to allow microbial colonization of substrates from the surrounding spring water. Control tubes were sealed with intact rubber bung caps and injected with 0.2 μm filtered spring water prior to incubation as a control for degradation resulting from the geochemical properties of the water.

For the time series incubation experiment, six paired experimental/control tubes each containing one PHB gyroid were suspended in one geothermal spring (Maruia). The incubation experiment lasted for six months (April 2023 – November 2023), with one pair of tubes removed each month.

For the multi-substrate incubation experiment, two tubes containing three PHB gyroids each (12 tubes), two tubes containing three PET gyroids each, two tubes containing three PLA gyroids each, and two control tubes each containing one gyroid of each polymer were suspended in four different geothermal springs within the TVZ (32 tubes total). We also incubated one PHB tube (three gyroids), one PET tube (three gyroids), one PLA tube (three gyroids), and one control tube with one gyroid of each polymer (4 tubes total) in Maruia spring in the Southern Alps. Tubes were left to incubate for up to three months (May 2023 – Sep 2023) before removal.

### Post incubation processing

The incubation chambers were removed from the springs, sealed with sterile parafilm, and transported on ice to the University of Canterbury for processing. In the laboratory, the spring water decanted and replaced with sterile PBS buffer (per L H_2_O: 8 g NaCl, 1.44 g Na_2_HPO_4_, 0.2 g KCl, 0.22 g KH_2_PO_4_). 5 ml PBS buffer was added to tubes with a single substrate (time series incubation), and 15 ml was added for tubes with multiple substrates (multi-polymer incubation). The re-filled tubes were then sonicated in a 37 kHz ultrasonic water bath (Elma Schmidbauer GmbH, Singen, DE) for an hour to remove the biomass from the substrates. After sonication, the substrates were removed from the PBS buffer, the tubes were centrifuged at 15,000 rpm for 5 min to concentrate the shaken biomass, the supernatant decanted, and the biomass was re-suspended in a new 2 ml aliquot of PBS buffer. The biomass-free substrates were then left to dry in a 55 °C oven for 48 h and were then re-massed to determine any loss (Dataset S1).

### DNA Extractions of biofilm

DNA was extracted from a 1 ml subsample of the resuspended biofilm from the 27 experimental incubation chambers from the multi-substrate incubation using the Qiagen DNeasy^®^ Powersoil^®^ Pro Kit (Qiagen, Germantown, MD, USA) following the manufacturer’s recommendations. Extraction blanks were included to test for any possible contamination introduced during the DNA extraction. Extractions were performed in a Class II biohazard hood to prevent contamination during the extraction process.

### Microbial analyses via marker gene sequencing

Sequencing was performed following the methods described in Dragone et al. (21). The DNA aliquots extracted from each of the 27 samples and their associated four extraction blanks were PCR-amplified using a primer set that targets the hypervariable V4 region of the archaeal and bacterial 16S rRNA gene (515F: 5’-GTGCCAGCMGCCGCGGTAA-3’ and 806-R: 5′-GGACTACHVGGGTWTCTAAT-3′) (22, 23) following the methods described in (24, 25). These primers included the appropriate Illumina adapters and unique 12 - bp barcode sequences to permit multiplexed sequencing (26). Nine no-template PCR blanks were included with each set of PCR amplifications to control for contamination. Amplifications were performed in duplicate on a SimpliAmp Thermal Cycler (Thermofisher Scientific, Waltham, MA, USA) using Platinum^TM^ II Hot-Start PCR Master Mix (2X) (Invitrogen, Carlsbad, CA, USA) in 25 µL reaction volumes with 4 µL template. Cycling parameters consisted of an initial denaturation step at 94 °C for 3 min, followed by 35 cycles of denaturation at 94 °C (45 s), annealing at 50 °C (60 s), extension at 70 °C (90 s), and a final extension step at 72 °C for 10 min. Duplicated amplified products were combined, cleaned, and normalized to equimolar concentrations using Sera-Mag SpeedBeads (27) and were sequenced on the Illumina MiSeq platform (Illumina, San Diego, CA, USA) using the V2 2 × 150 bp paired-end Illumina sequencing kit at the University of Colorado Boulder. The sequencing data was demultiplexed using idemp (https://github.com/yhwu/idemp).

The 16S rRNA gene sequences were processed using the DADA2 pipeline v.1.16 (28) with samples pooled at the inference step. More details of the specific parameters used can be found at https://github.com/fiererlab/dada2_fiererlab. Sequences were quality filtered and clustered into exact sequence variants (ASVs), with taxonomy determined using a naïve Bayesian classifier method (29) trained against the SILVA v.138.2 reference database (30, 31). A minimum bootstrapping threshold required to return a taxonomic classification of 50 % similarity was used for analysis. Prior to downstream analyses, ASVs associated with chloroplast, mitochondria, eukaryotes (24 ASVs total), those unassigned to the phylum level (45 ASVs), and singletons and doubletons (7 ASVs) were removed. Filtered ASV tables can be found in Dataset S2.

Additional community analyses were performed in R v.4.3.3 (32) using the packages ‘mctoolsr’ (https://github.com/leffj/mctoolsr/), ‘vegan’ (https://github.com/vegandevs/vegan), and phyloseq (33). Richness, the number of distinct prokaryotic ASVs, was calculated from the filtered 16S rRNA gene ASV tables using the ‘specnumber’ function.

### Preparation of powdered plastics for culturing

Powdered PET and PLA were prepared using a kitchen blender; injection moulding grade PET pellets (TechSoft Ltd, Wellington, NZ) and PLA pieces recycled from 3D printing support material (PLA+ 3D printing filament, eSun ltd, CH) were blended in bursts of 30 s to avoid overheating until a fine powder was obtained. PHB polymer was purchased in powder form (PHI003, NaturePlast, Mondeville, FR) and was used unmodified. All polymer powders were sieved to 100 – 300 µm.

### Cultivation

Putative plastic degrading microorganisms were selected through the production of a clearing zone in a plastic- amended thin overlayer on top of modified DSMZ Media 150a (see supplemental methods) prepared with noble agar (Thermo Fisher Scientific, Waltham, MA, USA) (34–36). Briefly, the medium was prepared by adjusting the pH of the DSMZ Media 150a to reflect the pH of the source spring (Figure 1, Dataset S1, S3 for details). The overlayer was then prepared by adding 10 g of one of the powdered plastic polymers (PET, PHB, PLA or a no-= additional control) to a 250 ml aliquot of pH adjusted base medium. All media was sterilized by autoclaving (121 °C; 15 PSI, 20 min) after which filter-sterilized chelated iron solution (3 ml/l), methanotroph trace element solution (3 ml/l), and methanogen trace element solutions (1 ml/l) was added to each medium aliquot. The base medium was then poured into 58 cm^2^ Petri dishes. After the base medium had solidified, 10 ml of the plastic-infused overlay was aseptically added on top of each base to create a thin, opaque layer of suspended plastic powder.

A 50 µl aliquot of the resuspended biofilm from gyroids from each growth tube from each hot spring was pipetted on five grow plates of each polymer type prepared for that spring. Plates were then incubated at either 45 °C, 55 °C, 65 °C, 70 °C, or 80 ^°^C depending on which was closest in temperature to the incubation spring (see Figure 1, Dataset S1, S3). Three uninoculated “blank” plates per plastic type per spring were incubated with the growth plates as controls for contamination. Plates were incubated for eight weeks under aerobic conditions and were checked weekly. Axenic cultures of all colonies that created clearing zones in the plastic overlayer (see Figure 4 for an example) were generated via three passages of single colonies on plastic-overlay solid media. Additional culturing information can be found in Dataset S3.

### Whole genome sequencing and identification of plastic degrading isolates

DNA from the two isolated colonies that formed clearing zones was extracted using a DNeasy PowerSoil Pro Kit (Qiagen, Hilden, Germany) with partial automation on a QIAcube Connect (Qiagen) following the manufacturer’s instructions.

Whole genome sequencing of gDNA extracted from the two isolates was performed by SeqCenter (Pittsburgh, PA, US) following the sequencing provider’s standard protocol. Briefly, Illumina sequencing libraries were prepared with the Nextera DNA Flex library preparation kit for a target insert size of 280 bp (Illumina, San Diego, CA, US). DNA was sequenced on an Illumina NovaSeq X Plus sequencer in multiplexed runs with a 2 × 151 bp paired end-read chemistry. Demultiplexing was performed with bcl-convert v. 4.2.4 (Illumina, San Diego, CA, US).

The raw paired-end sequencing reads were assessed for quality with FastQC v.0.12.0 (https://github.com/s-andrews/FastQC). Illumina adapters were removed, and quality filtering was performed with Trimmomatic v.0.39 (37). Genome assembly was performed with Unicycler v.0.5.1 (38) and completeness and contamination was assessed with CheckM v.1.1.6 (39). Once a high-quality genome was assembled (defined as >95 % complete, <5 % contamination), open reading frames (ORFs) were predicted with Prodigal v.2.6.3 (40), and taxonomy was assigned using the classify workflow of GTDBtk v.2.4.0 (41) and confirmed with the de novo workflow of GTDBtk, which used FastTree (42) to place our genomes within the GTDB-tk reference bac120 tree (43) based on multiple sequence alignments (MSA) of amino acid sequences. To further confirm the taxonomic identifications of the two plastic degraders provided by GTDBtk, we used phyloFlash v.3.4.2 (44) to extract the 16S rRNA SSU genes from the genome sequences and assign taxonomy based on the SILVA database v.138 (30, 31, 45). Trees were visualized and annotated using iTOL v.7.2 (46).

Identification of PHA metabolism genes in the two genomes was performed by comparing the translated amino acid sequences against references sequences from seven different databases related to PHA/PHB degradation and/or synthesis with the blastp function of DIAMOND v.2.0.11 following parameters described in (24, 47) (query coverage: 80 %, e-value threshold: 10^-10^). Databases include: PlasticDB (56 reference sequences) (5), PMBD (16 reference sequences) (48), PAZY (13 reference sequences) (49), COG (503 reference sequences) (50), KEGG (542 reference sequences) (51), the Gene Ontology Resource (7 reference sequences) (52, 53), and Uniprot (36 reference sequences of PHA/PHB depolymerase restricted to Swiss-Prot reviewed entries) (54). As described in Dragone et al. (47), we used a lower percent identity (>60 %) to match our genomes to the reference due to of the small number of reference sequences available for plastic degrading genes and the limited phylogenetic diversity from which they were collected (5). Matches above this identity threshold were considered positive identifications. Whole genome sequencing data can be found in Dataset S3.

### Predictive modeling of thermophilic plastic degraders

Details regarding the sampling process and characterization of the geothermal spring samples used for the biogeography analysis can be found in Power et al. (16). To summarize, they collected samples from >900 geothermal springs across New Zealand (http://1000Springs.org.nz), measured 46 physicochemical parameters of the samples, and sequenced extracted DNA with an Ion PGM System using the Earth Microbiome primer pair F515/R806, described previously. We accessed the raw 16S rRNA gene sequences from the European Nucleotide Archive under accession number PRJEB24353. The 16S rRNA gene sequences were processed using the DADA2 pipeline v.1.16 (28), described previously, but with the recommended parameters for processing Ion Torrent Data. Prior to downstream analyses, ASVs associated with chloroplast, mitochondria, eukaryotes (939 ASVs total), those unassigned to the phylum level (2303 ASVs), and singletons and doubletons (2720 ASVs) were removed. Samples were only used for downstream analysis if >1000 reads were recovered from them (899 samples).

To add to the two 16S rRNA gene sequences recovered from our thermophilic plastic-degraders, we searched in PlasticDB v.01-27-2025 for all bacterial taxa previously reported in literature to degrade plastic polymers at temperatures >50 °C (5). We then used the PlasticDB record for each hit to search for the 16S rRNA gene sequences of the specific culture listed in the entry (Dataset S3). If the specific 16S rRNA gene sequence of a culture could not be found, but the taxonomic classification listed a species and strain, we used the 16S rRNA gene sequence of the associated type strain. If a PlasticDB entry was not classified to a unique species (i.e. “*Streptomyces* sp.”) and we could not find a 16S rRNA sequence associated with that specific culture, it was excluded. The 16S rRNA gene sequences recovered from thermophilic plastic degraders (56 from PlasticDB, 2 from our isolates) were matched against the 16S rRNA gene sequences associated with each ASV recovered from the 1000 Springs Project dataset (described previously, Dataset S4) using VSEARCH v2.22.1 (--strand both --notrunclabels --iddef 0 --id 0.97 --maxrejects 100 -- maxaccepts 100) (55). Putative thermophilic plastic-degraders were considered as present in a sample if any ASV that matched to our references sequences above our stated threshold was present and the proportion of thermophilic plastic-degraders was based on the sum of reads from these matched ASVs (Dataset S4).

To determine what environmental and geochemical variables might best predict which of the 899 geothermal springs would have thermophilic plastic degraders, we used the R package ‘rfPermute’ and performed a random forest analysis with 100 trees and 3 variables tried at each split to identify the most important predictors. The factors used in our models were chosen from a total of 29 different measurements taken from the 46 reported by Power et al. (16). Variables that were not measured in at least 75 % of the samples were not included in the analysis. Variables were considered predictors of the presence of thermophilic plastic degraders so long as that variable increased the MSE by at least 4 % (*p* < 0.05). To further explain these relationships, the abundance of the sum reads of all matched ASVs were plotted against the respective predictive variable and generalized additive models (GAMs) were used to predict under what conditions abundance was maximized. We used the gam.check function of the r package ‘mgcv’ (56) to calculate diagnostics for each model. Parameters, including k’, effective degrees of freedom, and k-index can be found in Dataset S4.

### Additional plotting and analyses

Statistical tests were performed using base R functions ‘wilcox.test’ and the packages ‘rstatix’ (https://github.com/kassambara/rstatix). Plotting was performed using the R packages ‘ggplot2,’ ‘cowplot’, and ‘mctoolsr’. The map in Figure 1 was created in QGIS v.3.44 (57) with the "QuickMapServices" plugin using the Carto Positron basemap (58).

## Results

### Time series incubation

Average pH measured at Maruia spring across the six-month incubation was 7.97 (range: pH 7.77 – 8.30), average temperature was 57.0 °C (52.5 – 61.2 °C), and average conductivity was 1116 mS (range: 910 – 1340 mS). The average mass lost by our experimental PHB gyroid setup after a six-month incubation in Maruia spring was 8.5 % (w/w; mass loss × g). The control PHB gyroid lost 5.4 % (w/w; mass loss × g) of mass during the same incubation. Approximate rates of plastic degradation across the experiment for the experimental gyroid was 0.036 % · day^-1^ and 0.026 % · day^-1^ for the control gyroid (Figure 3).

**Figure 3:**
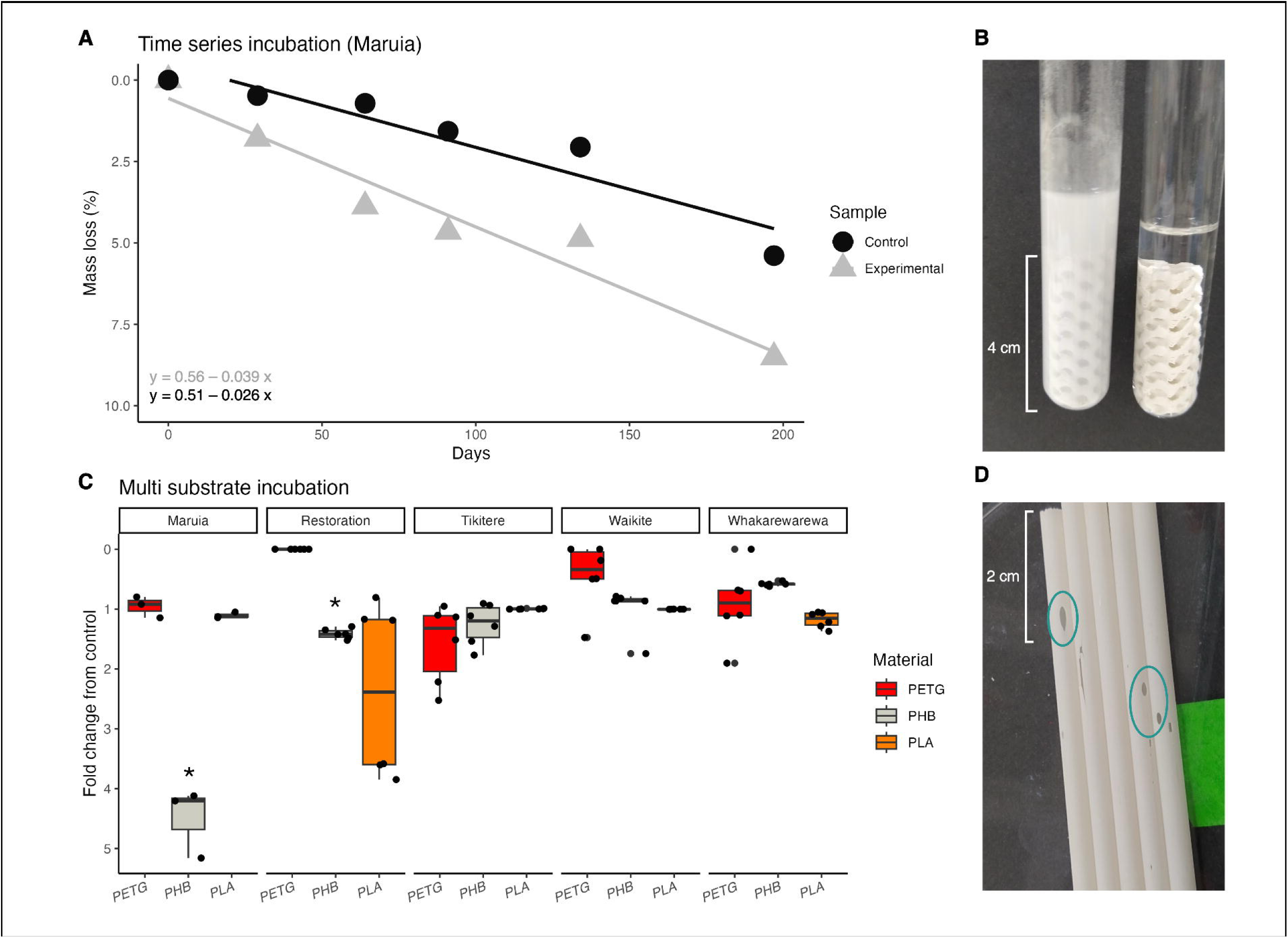
Mass loss experimental results. **A)** The results of the time series incubation of our three polymers: Polyethylene terephthalate (PET), Polylactic acid (PLA), and Polyhydroxybutyrate (PHB). Mass loss for the control and experimental gyroid are displayed over the 6-month incubation. A line of best fit, predicted using a generalized linear model method is included for each treatment, with equations listed at the bottom left. **B)** Difference between the experimental gyroid (left) compared to the control gyroid (right) after incubation and sonication to remove biofilm. **C)** The results of the multi substrate incubation. Box plot displays fold change from average mass loss across control gyroids (n = 2) for each experimental gyroid (n = 6) after three months. For the Maruia incubation only, a single experimental chamber was collected at the three-month time point (n = 1 control gyroid, n = 3 experimental gyroids). Results are grouped by spring and colored by polymer type. Springs/polymers from which plastic degrading organisms were isolated are indicated with a star (*). For more information on how plastic degradation was determined, see Methods. **D)** Evidence of microbial plastic degradation on PHB-H straws (circled in green) incubated for three months in Maruia spring concurrently with the multi-substrate incubation.

### Multi-substrate incubation

The measured temperature, pH, and conductivity of each spring at the start and end of the incubation experiment can be found in Dataset S1. After the three-month incubation, average % loss (w/w; mass loss × g) of PHB gyroids was 23.1 % (2.7 – 67.7 %), was 0.71 % (0 – 3.3 %) for PETG gyroids, and 62.4 % (3.7 – 94.8 %) for PLA gyroids (Figure 3). Control gyroids lost on average (% w/w; mass loss × g) 21.8 (0.66 – 38.8 %) for PHB, 0.83 (0 – 2.9 %) for PETG, and 56.8 (3.5 – 95.0 %) for PLA. Gyroid mass before and after incubation and fold change from control gyroids can be found in Dataset S1.

### Cultivation-independent substrate comparisons

After the three-month incubation, visible biomass could be observed in all experimental gyroids exposed to the spring but not on the sealed control gyroids (see Figure S2 as an example). DNA was extracted from biomass removed from the gyroids in each of the 27 experimental chambers used in our multi-substrate incubation.

The sequences recovered from the blank did not suggest signs of contamination. Just 35 reads from nine ASVs were recovered from the extraction blanks while 37 reads from nine ASVs were recovered from the no template controls. In total, the ASVs assigned to these samples were neither the most abundant (average <0.1 % of total reads assigned to samples), nor are they typical laboratory contaminants (59).

All 27 of the biofilm samples yielded enough PCR amplifiable DNA to characterize the microbial communities using marker gene sequencing (Dataset S2). 16S rRNA gene reads averaged 5406.5 reads sample^-1^ (12 – 30579 reads) with an average richness of 75.2 ASVs sample^-1^ (5 – 345 ASVs) (Figure S3). Of the 594 unique ASVs identified, 538 ASVs were bacterial and 56 ASV archaeal. A total of 59 bacterial phyla were detected with the five most abundant being Pseudomonadota (24.3 % of total reads), Thermoproteota (16.3 %), Thermoplasmatota (8.3 %), Bacillota (8.3 %), Methanobacteriota (4.5 %) (Figure 4). The ten most abundant ASVs are presented in Table S1 and include the families Methanothermobacteriaceae (2 ASVs), Spirochaetaceae (2 ASVs), Thermodesulfovibrionaceae, Methanosaetaceae, Syntrophomonadaceae, and unidentified families of the orders Veillonellales-Selenomonadales, Thermoanaerobacterales, and Acetothermiia; KB1 group. The complete ASV table can be found in Dataset S2.

The spring in which the gyroid was incubated was the strongest predictor of observed variation in prokaryotic community dissimilarity across the 27 samples (PERMANOVA, r^2^ = 0.39, *p* = 0.001, Figure 4). All springs varied significantly in their community composition (*p* < 0.05). Of the physicochemical parameters measured at each spring, pH was the only variable identified as a significant driver of this dissimilarity (MRM, *p* = 0.02; bioenv: r^2^ = 0.52). Biofilms from springs with low pH (<7, Whakarewarewa and Tikitere) had an average richness of 14.4 ASVs (5–32) and were dominated by Pseudomonadota (33.6 %), Thermoproteota (20.0 %), and Thermoplasmatota (17.8 %) with all other phyla making up <3 % of the total reads. Six of the top 10 ASVs in the more acidic springs were Archaea (Table S1), which made up 75.8 % of the reads recovered from these samples. Springs with neutral to moderately alkaline pHs (>7) had an average richness of 123.8 ASVs (28 – 345 ASVs) with the most abundant phyla in these springs being Pseudomonadota (16.9 %), Thermoproteota (13.6 %), Bacillota (12.1 %), Methanobacteriota (7.7 %) and Aquificota (5.1 %). Just three of the top 10 most abundant ASVs were Archaea (Table S1), with Archaea making up just under a quarter (23.1 %) of reads on average.

Substrate type also explained some of the variation in microbial community composition (PERMANOVA, r^2^ = 0.07, *p* = 0.027). Most (370) ASVs were identified on at least two of the different substrate types. Sixty ASVs were only found in PET biofilms, 23 were only found in PLA biofilms, and 141 were only found in PHB biofilms (Figure S4). 63 ASVs were identified as being significant drivers of the dissimilarity between the three substrates (SIMPER, *p* < 0.05, Figure 4). Of these, the most abundant genera that were consistently enriched in biofilms on PET included *Methanothermobacter* (6.1 % of reads from PET samples on average), *Fervidicoccus* (5.0 %), *Caldimicrobium* (4.3 %), *Methanothrix* (11.8 %) and an unidentified Bathyarchaeia (1.6 %) (Figure 4). The most abundant genera associated with PHB biofilms include *Herminiimonas* (2.2 % of reads from PHB samples on average), *Chloroflexus* (1.6 %), *Chlorogloeopsis* (0.73 %), *Thermus* (0.59 %), and *Fervidicoccus* (0.54 %) (Figure 3). The most abundant genera associated with PLA biofilms included *Citrobacter* (14.4 % of reads from PLA samples on average), *Afipia* (1.8 %), *Chloroflexus* (0.93 %), *Candidatus* Nicrosocaldus (0.90 %), and *Lysinibacillus* (0.69 %) (Figure 4). Indicator taxa analysis (multipatt) identified ASV 24 (Archaea; Thermoproteota; Nitrososphaeria; SCGC AB-179-E04; NA; NA) as indicative of PET biofilms and ASV 181 (Bacteria; Pseudomonadota; Gammaproteobacteria; Enterobacterales; Enterobacteriaceae; *Citrobacter*) as indicative of PLA biofilms. No PHB indicator taxa were identified. Taxa previously been reported to be plastic-degraders are listed in Table S2

### Enrichment, isolation, and characterization plastic-degrading taxa

No growth was observed on any of the control plates after eight weeks, nor did any colonies grow on plates inoculated from gyroids incubated in Whakarewarewa and Tikitere springs. No clearing zones were noted on any PET or PLA screening plates from any spring, nor from any PHB plates from Waikite Valley after eight weeks’ incubation.

PHB-degrading colonies (i.e. colonies that formed clearing zones on the PHB screening plates) were identified on two solid medium plates within three and 14 days of initial inoculation (Dataset S3). The single colony morphotype recovered from the PHB biofilm from the Maruia Spring incubations (PHB tube 1) degraded PHB at 55 °C, while the other colony morphotype was from Waikite Restoration PHB biofilm (PHB tube 5) and degraded PHB at 45 °C. High quality genomes were assembled from both isolates (Dataset S3). The Maruia PHB-degrading isolate was identified as a *Rubrobacter* sp. [Maruia] (phylum Actinomycetota; 99.26 % ANI, 95 % AF to GTDB accession GCF_028617085.1, Figure S5) and the Waikite Restoration PHB degrading isolate was identified as a *Cupriavidus* sp. [WR] (phylum Pseudomonadota; 98.7 % ANI, 90 % AF to GTDB accession GCF_008632125.1, Figure S6). Taxonomic identification was confirmed through 16S rRNA gene sequence similarity and average nucleotide identity.

Both genomes were found to contain genes that matched references sequences of previously described PHA metabolism pathways (Figure 4, Figure S7, Dataset S3). *Cupriavidus* sp. [WR] matched to references sequences from the COG database (1 match), Uniprot (1), PlasticDB (3 matches), and PAZY (4). These included an extracellular PHA depolymerase (PhaZ6), intracellular PHA depolymerases (PhaZ1, PhaZ2), 3-hydroxybutyrate-oligomer hydrolase genes (PhaY1, PhaY2), and a class I poly(R)-hydroxyalkanoic acid synthase gene (PhaC1). *Rubrobacter* sp. [Maruia] only matched to one reference sequence, a class III poly(R)-hydroxyalkanoic acid synthase (PhaC3) from the COG database.

### Predictive modeling of thermophilic plastic-degraders

We recovered 56 16S rRNA gene sequences from the 193 entries associated with thermophilic plastic degraders in PlasticDB (5). These sequences, plus the two 16S rRNA gene sequences recovered from our isolates, matched to 246 ASVs recovered from the reprocessed sequence data from Power et al. (16). Reads assigned to these ASVs were identified in 429 of the 898 samples (47.4 % of samples). The average abundance of database-matched ASVs was 13.5 % of total reads per sample (0 – 41.1 % of reads).

One variable, spring pH (% IncMSE = 6.97, *p* = 0.01) was identified as an important predictor of the abundance of thermophilic plastic degraders in a spring. Aluminum concentration was also identified as a significant driver (*p* = 0.04) but did not increase the mean standard error by at least 5 % (Table S3). Matched ASVs were identified in samples ranging in pH from 1.34 – 9.7 and with an aluminum concentration from 0.18 – 198.7 mg/L. Predicted maximum abundance of matched ASVs was between pH 1- 2.3 (Fig. 4) and at an aluminum concentration of 6.5 – 65 mg/L (Figure S8).

## Discussion

To meet our objectives, we incubated 120 plastic substrates (87 experimental gyroids, 33 control gyroids) of three different polymers (PET, PLA, PHB) in five springs across New Zealand (four in the TVZ, and one in the Southern Alps) for up to 6 months to observe any relationships between geothermal spring microorganisms and plastic degradation. These springs varied in temperature from <45 – 85 °C and in pH from <5 – 9. Using a combination of cultivation-independent marker gene sequencing, targeted cultivation paired with whole genome sequencing, and predictive modeling, we were able to show how introduced plastic changes the microbial communities in geothermal springs, identify and grow thermophilic microorganisms capable of degrading certain plastic polymers, and predict which springs may contain more thermophilic plastic-degraders.

### PHB degrades in geothermal springs at rates above that of the associated sterile controls

The majority of our gyroids, including the controls, reduced in mass over the course of the incubations. In the time series incubation, the PHB control gyroids lost mass at an approximate rate of 0.52 % (w/w) per month (Figure 3) and was most likely the result of chemical and physical degradation of polymers and/or fillers induced by spring water. We expect that the experimental gyroids experienced a similar mass loss due to physical/chemical degradation, but that the increased degradation rate compared to the control was the result of microbial activity. For this reason, we focused on the magnitude of change from the mass lost from the control (i.e. fold change) rather than the absolute loss. However, what we describe below represents inferred, rather than directly demonstrated, microbially mediated plastic degradation.

Both of our incubation experiments suggests that microorganisms in geothermal springs contribute to the degradation of introduced PHB: In the time series incubation, the experimental gyroids lost mass at a rate 1.5 × faster than the controls (Figure 3) and for the multi-substrate incubations, potential microbial degradation (fold change from control >1) of PHB was detected in three of our springs with the largest mass loss compared to control measured in Maruia spring (temp: 55 °C; avg fold change: 4.5, Figure 3).

PHB is the most common form of polyhydroxyalkanoate (PHA) – a class of polyester produced through the activity of microorganisms (60) that consist of chains of hydroxyalkanoate groups linked together by ester functional groups (61). Microbially produced plastics like PHB have historically been associated with carbon storage (62) and produced intracellularly as water - insoluble inclusions, or granules when carbon is in excess (as described in refs (63–65)). If carbon becomes limited, these reserves can be broken down and used for energy (66). Given that PHB is known to be readily degraded by mesophilic and extremophilic organisms in a variety of copiotrophic and oligotrophic systems, this finding is not surprising (5, 66–68). However, we note that PHB degradation in our springs (∼8 % loss after 6 months) was much less than what has been measured in other aquatic systems (over 25 % loss in sea water in as little as one month) (69, 70).

### Microbial degradation of PET and PLA could be a result of metabolism of fillers/additives

Of our three polymers, PET gyroids were the most resistant to microbial degradation (Figure 3). This is not a surprise; while some PET-degrading microorganisms have been isolated (see refs (5, 71) for examples), PET breakdown is energy intensive and not typically favorable if alternative energy sources are available (72, 73). At higher temperatures (>65 °C) and more acidic pH, PET becomes more flexible and therefore more biodegradable if capable organisms are present (73, 74). This may explain why the only signs of mass loss by PET that could be attributed to microbial activity occurred in one of our highest temperature springs (Tikitere, Figure 3).

PLA gyroids did appear to lose mass as a result of microbial activity in three springs (Maruia, Waikite Restoration, and Whakarewarewa). PLA, a bioplastic produced by organic sources, has been touted as a “green” plastic alternative to petroleum-based polymers. However, PLA is still resistant to biodegradation and can take several years to break down outside of industrial compost facilities (75, 76). Our results suggest that geothermal springs may be a source environment to isolate more effective PLA degrading organisms, especially since microbial degradation of PLA has been shown to increase under thermophilic conditions (>50 °C) where polymer chains are more flexible and susceptible to enzymatic attack (76, 77). However, we saw no evidence of microbial plastic degradation at temperatures >70 °C, which might suggest an upper temperature limit on microbial PLA degradation under thermophilic conditions.

While our incubation results support (14) hypothesis that Aotearoa New Zealand’s hot springs contain microorganisms capable of degrading plastics, we cannot say whether the mass loss by our PET and PLA gyroids are a result of microbial degradation of the polymer or of plastic additives. The 3D-printing stock used to make our PETG and PLA gyroids contained plasticizers, stabilizers, and colorants (see methods for specifics) (78). The mass lost attributed to microbial activity could be the result of microbial breakdown of these accessory compounds instead of the polymers (as described in (79, 80)). For example, the PLA filaments contained calcium carbonate, which could either act as a source of carbon and/or potentially dissolve in highly acidic pools. Future work should account for this possibility and use raw polymers for the collection of the biofilms. However, we also saw evidence that additives may be released from introduced plastics just through exposure to the *in situ* abiotic conditions experienced in our springs. For example, sterile PLA gyroids in our highest temperature springs not only lost significant mass (>85 % w/w mass, Figure 3) but also lost color to the surrounding water of the chamber (Figure S2). Future work should determine whether the release of these chemicals at high temperatures has any impact on the chemical properties of these fragile environments and their associated biological communities.

### Introduced plastics can influence microbial community composition

Observed differences in the composition of biofilm communities on our incubated substrates were largely explained by the physicochemical conditions of the spring in which the substrate was incubated in (Figure 4), with difference in pH identified as the variable contributing the most to this dissimilarity. More specifically, biofilms from substrates incubated in more alkaline pH springs (>7) had more diverse communities dominated by Bacteria (Figure 4, Table S1). Biofilms from substrates incubated more acidic springs (<7) were less diverse and were dominated by Archaea (Figure 4, Table S1). Previous work identified similar drivers of microbial communities within the TVZ across a greater number of springs (16)

A smaller proportion of biofilm community dissimilarity was explained by the polymer type. While the majority of the ASVs identified across our study were shared across two or more polymers, ASVs assigned to ∼50 genera were identified as significant drivers of dissimilarity between biofilm microbial communities across polymer types. The most abundant genera associated with each polymer are known biofilm formers (i.e. *Methanothermobacter –* PET, *Caldimicrobium –* PET*, Herminiimonas –* PHB*, Citrobacter –* PLA*, Chloroflexus –* PHB) (81–85). However, just four genera (*Citrobacter, Lysinibacillus, Corynebacterium,* and *Pseudomonas)* has previously been reported to degrade any plastic polymer, with just *Pseudomonas* reported to degrade any of the three polymers used in this study (Table S2) (5). These results suggest that plastics introduced into geothermal systems do not enrich for organisms capable of using the polymer as an energy source (as has been described in other systems (86)). Instead, introduced plastic mainly appears to act as a surface for biofilm formers to adhere to, similar to what has been described in marine systems (87). However, we have yet to fully understand the plastic-degrading capability of the microbial world and do not know the full diversity of organisms that degrade plastic (4, 88). It is likely that some of these organisms associated with communities on each polymer have undescribed plastic degradation capabilities.

### The role of PHB degradation by thermophiles

Two of our isolates formed clearing zones in the PHB screening medium, confirming plastic-degrading capabilities. One isolate, *Rubrobacter* sp. [Maruia], was confirmed to be a thermophilic plastic degrader that degraded plastic at 55 °C (Figure 5). The other isolate, *Cupriavidus* sp. [WR], degraded plastic at 45 °C. *Cupriavidus* has previously been identified as a plastic-degrading organism (based on the PlasticDB database (5)). To our knowledge, no reports exist of *Rubrobacter* degrading plastic (5).

**Figure 4:**
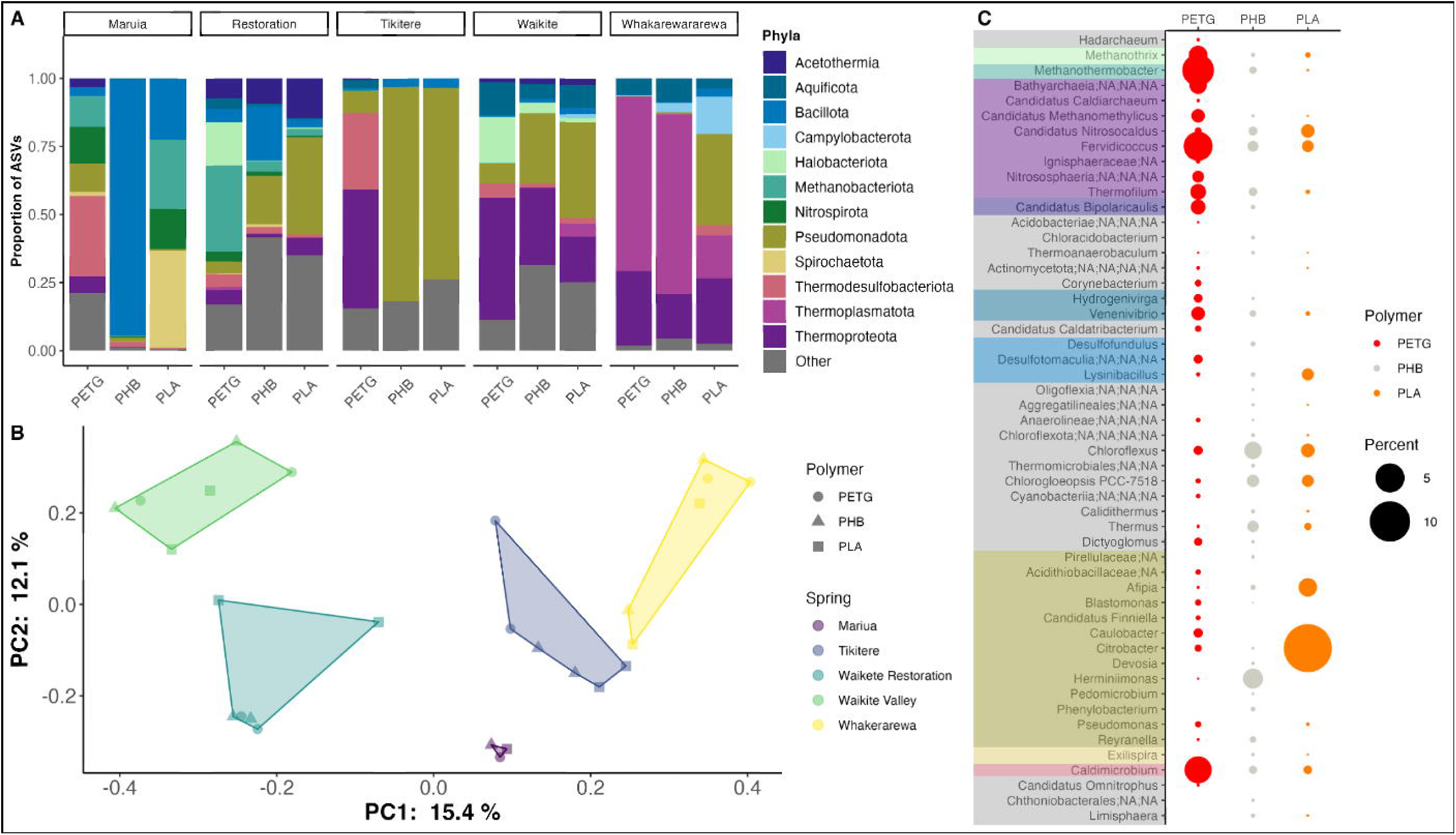
Biofilm comparisons across site and polymer. **A)** Stacked bar chart of the 12 most abundant bacterial and archaeal phyla across the multi substrate incubation. Relative abundance displayed the combined relative abundance from the two replicates for each polymer type at each spring. **B)** Principal Coordinates Analysis (PCoA) displaying the Bray Curtis dissimilarity across the different springs. Shape designates the polymer type while the color displays the spring that the sample was incubated in. **C)** Percent of ASVs associated with genera identified through indicator taxa analysis (Multipatt and SIMPER) to be driving the differences in microbial communities between the three polymer types across all springs. Genera are colored based on the phyla displayed in A.

**Figure 5:**
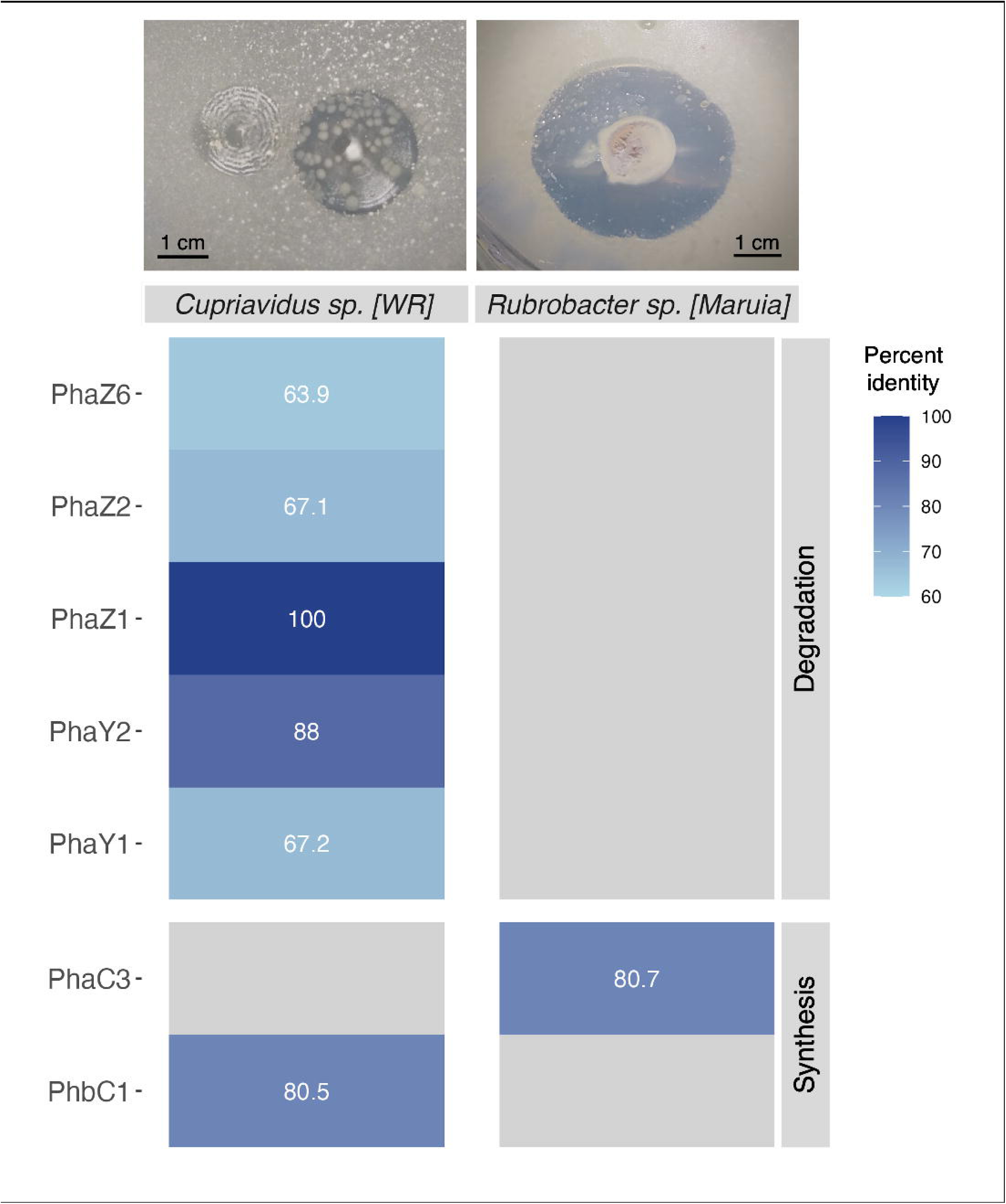
Summary of the PHA/PHB metabolism genes identified in the assembled genomes of our two isolates. Genes are grouped into “degradation” and “synthesis.” Heatmap intensity displays the percent identity match of the gene found in our isolate to the reference gene sequence. Photos of the clearing zones formed by each isolate in the screening media are displayed above the heat map.

We were able to reconstruct near-complete high quality draft genomes from our two PHB degraders (>97 % complete, <2.5 % contamination).The genes identified in the reconstructed genomes of our *Cupriavidus* sp. [WR] isolate supports the hypothesis that certain microbes are scavenging extracellular PHB to use as source of carbon and energy, a strategy which has been described in other geothermal environments (11, 68). Our *Cupriavidus* sp. [WR] isolate encoded an extracellular PHA/PHB depolymerase gene (PhaZ6, Figure 5) which putatively would enable the organism to break down scavenged PHB into oligomers and monomers (49, 89). Moreover, we also identified PhaY1 and PhaY2 genes, used by organisms to further metabolize these oligomers (49, 90).

While no genes associated with any PHB degradation pathway were identified in the genome of *Rubrobacter* sp. [Maruia], we expect it contains a similar extracellular PHB degradation pathway, given the completeness of the genome and that we observed clear extracellular PHB degradation on our screening medium (Figure 5). Most plastic degradation genes have not been well characterized in more than one organism (Chow et al. 2023, Buchholz et al. 2022). While it is possible that our *Rubrobacter* sp. is utilizing a previously undescribed PHB degradation pathway, it is more likely that no genes have been described in closely related organisms (unlike our *Cupriavidus* isolate encodes PHB- degrading gene with high homology to previously characterized PHB degradation genes within the genera (91)).

While we were not able to grow any organisms that would directly degrade PET or PLA in our assays, further testing and isolation campaigns may reveal this capability in other organisms and springs. Our cultivation methods were narrow, relying on just one base medium type, one set of laboratory conditions, all under aerobic conditions. We expect that a larger cultivation campaign, targeting a larger number of springs, and using a wider variety of media types, plastic polymers, and growth conditions may successfully capture organisms capable of degrading PET, PLA, or other polymer types.

### New Zealand’s hot springs are a potential source of novel microbially produced plastics

Although we did not directly test this capability, genomic evidence indicates that both isolates may be able to synthesize and degrade intracellular PHB. Each genome contained high-identity matches (>80 %) to intracellular PHA synthase genes (PhaC)—Type I in *Cupriavidus* sp. [WR] and Type III in *Rubrobacter* sp. [Maruia] (Figure 5). In addition, *Cupriavidus* sp. [WR] encoded the intracellular PHB depolymerases PhaZ1 and PhaZ2 (Figure 5). These findings suggest that both organisms may produce PHB as an intracellular carbon reserve that can be mobilized during nutrient limitation, consistent with observations in other systems (65, 68, 90)

Thermophilic PHB--producing organisms offer many of the same advantages as thermophilic plastic degraders, including faster production, greater efficiency, and reduced risk of contamination (92, 93). Notably, our results suggest that geothermal organisms from New Zealand—such as the two isolates described here—may be capable of generating more thermostable PHBs. Such materials could facilitate broader adoption of PHB as a sustainable alternative to petroleum-derived- plastics like polyethylene and polypropylene (60).

### New Zealand hot springs support a range of putative plastic degraders

We leveraged the extensive geochemical, environmental, and microbial analysis performed by Powers et al. (2018) to identify springs with a high potential for harboring additional plastic degrading taxa beyond the five springs investigated in this study. By comparing the 16S rRNA gene sequences recovered from known thermophilic plastic degraders (including our two isolates) against the marker gene sequencing data created by the ‘1000 Springs Project dataset,’ we were able to identify pH as the most important predictor of thermophilic plastic degraders, with the greatest abundances of thermophilic plastic degraders predicted in springs with acidic pH (from 1 – 2.3, Figure S8) (See Dataset S4 for more details).

Acidic conditions have previously been linked to degradation of plastic. Ariza-Tarzona et al. (2020) found that low pH environments promoted polymer breakdown of polyethylene (94), which could then make the polyethylene easier for microorganisms to degrade (73). In fact, early work into plastic degradation by extremophilic organisms focused on acidophilic taxa specifically because of the ‘softening’ effect of acids on plastics (9) – an effect that is likely compounded at high temperatures (7, 8, 11). However, we note that predicted thermophilic plastic-degraders were identified across the pH range encompassed within the TVZ. Given the limited taxonomy of known organisms that can degrade plastic at temperatures >50 °C (9, 11), we believe that future studies should focus on springs across the full pH range, not just within our predicted maximum range. For example, (14) identified springs with moderate thermophilic conditions (30-40 °C) as containing higher relative abundances of putative plastic-degrading taxa.

### Conclusions

Thermophilic plastic-degrading microorganisms are present in the geothermal springs of Aotearoa New Zealand. Moreover, these organisms appear capable of degrading certain plastics introduced into these environments. Microbially produced polymers and biopolymers appear to be the most susceptible to microbial degradation by geothermal spring organisms, while petroleum-based polymers are not degraded easily. While the pathways identified in our plastic-degrading isolates suggest they may be able to metabolize and even produce plastics for carbon, the primary use of plastics by geothermal microbial communities does not appear to be as an energy source. Instead, they appear to mainly serve as surfaces for biofilm formation, composed primarily of non-plastic-degrading taxa. Not only do our findings support previous suggestions that New Zealand’s geothermal springs may be a source of novel plastic degraders, but they also suggest that they could be a reservoir of organisms capable of producing more thermostable, microbially derived plastics with improved performance characteristics.

## Acknowledgements

We wish to thank Prof. Noah Fierer, Jordan Galletta, Caihong Vanderburgh, and the Fierer Laboratory at CU Boulder for providing advice and material; the Stott Laboratory (XPHILES) and Craig Galilee at the University of Canterbury for laboratory assistance and technical support. We also acknowledge SeqCenter (Pittsburgh, PA USA) and the Center for Microbial Exploration at the University of Colorado Boulder (Boulder, CO USA) for sequencing services. We thank Maruia Hot Springs, Whakerewarewa – The Living Māori Village, Hell’s Gate (Tikitere), and Waikete Valley Hot Pools for providing us access to our incubation sites and mana whenua for their help. Samples collected from the Waikite restoration site via a New Zealand Department of Conservation permit.

We also acknowledge that these geothermal environments, and the microorganisms associated with them, are culturally significance to the Māori (the Indigenous People of Aotearoa New Zealand) and can be considered taonga (treasure). We thank Ngāti Tahu-Ngāti Whaoa, Ngāti Waewae, Ngāti Rangiteaorere, and Tūhourangi Ngāti Wāhiao who hold kaitiakitanga (guardianship) and are mana whenua of these sites and the microorganisms that reside there.

## Data Availability

All data used in this study are available in the main text or the supplementary material. The demultiplexed 16S rRNA gene sequencing data we generated from the gyroids, as well as the whole genome sequencing data from our two isolates are available in the National Center for Biotechnology Information Sequence Read Archive: BioProject ID PRJNA1448043

## Funding

This project was funded by the New Zealand Institute for Public Health and Forensic Science (PHF Science)’s Strategic Science Investment Funding from the Ministry of Business, Innovation and Employment, New Zealand [grant number C03 × 1701], under the project “Ōhanga Āmiomio—Tikanga led modelling of reduction and resource recovery from human waste and wastewater.” Financial support was provided to N.B.D. for this project by the Fulbright U.S. Student Program, which is sponsored by the U.S. Department of State and the New Zealand Fulbright Commission. The contents of this publication are solely the responsibility of the authors and do not necessarily represent the official views of the Fulbright Program, the Government of the United States, or Fulbright New Zealand.

## Notes

### Competing Interest Statement

The authors have declared no competing interest.

## References

1. Geyer R, Jambeck JR, Law KL. 2017. Production, use, and fate of all plastics ever made. Sci Adv 3:e1700782

2. Zalasiewicz J, Waters CN, Ivar do Sul JA, Corcoran PL, Barnosky AD, Cearreta A, Edgeworth M, Gałuszka A, Jeandel C, Leinfelder R, McNeill JR, Steffen W, Summerhayes C, Wagreich M, Williams M, Wolfe AP, Yonan Y. 2016. The geological cycle of plastics and their use as a stratigraphic indicator of the Anthropocene. Anthropocene 13:4–17

3. Horton AA, Walton A, Spurgeon DJ, Lahive E, Svendsen C. 2017. Microplastics in freshwater and terrestrial environments: evaluating the current understanding to identify the knowledge gaps and future research priorities. Sci Total Environ 586:127–141.

4. Shilpa, Basak N, Meena SS. 2022. Microbial biodegradation of plastics: challenges, opportunities, and a critical perspective. Front Environ Sci Eng 16:161.

5. Gambarini V, Pantos O, Kingsbury JM, Weaver L, Handley KM, Lear G. 2022. PlasticDB: a database of microorganisms and proteins linked to plastic biodegradation. Database 2022:baac008.

6. Bertocchini F, Arias CF. 2023. Why have we not yet solved the challenge of plastic degradation by biological means? PLoS Biol 21:e3001979.

7. James-Pearson LF, Dudley KJ, Te’o VSJ, Patel BKC. 2023. A hot topic: thermophilic plastic biodegradation. Trends Biotechnol 41:1117–1126.

8. Valdez-Nuñez LF, Rivera-Jacinto MA. 2024. Thermophilic bacteria from Peruvian hot springs with high potential application in environmental biotechnology. Environ Technol 45:1420–1435.

9. Pham VHT, Kim J, Chang S. 2024. A valuable source of promising extremophiles in microbial plastic degradation. Polymers 16:2109.

10. Roudaut G, Simatos D, Champion D, Contreras-Lopez E, Le Meste M. 2004. Molecular mobility around the glass transition temperature: a mini review. Innov Food Sci Emerg Technol 5:127–134.

11. Atanasova N, Stoitsova S, Paunova-Krasteva T, Kambourova M. 2021. Plastic degradation by extremophilic bacteria. Int J Mol Sci 22:5610.

12. Trego A, Palmeiro-Sánchez T, Graham A, Ijaz UZ, O’Flaherty V. 2024. First evidence for temperature’s influence on the enrichment, assembly, and activity of polyhydroxyalkanoate-synthesizing mixed microbial communities. Front Syst Biol 4:1375472.

13. Estévez-Alonso Á, Pei R, van Loosdrecht MCM, Kleerebezem R, Werker A. 2021. Scaling-up microbial community-based polyhydroxyalkanoate production: status and challenges. Bioresour Technol 327:124790.

14. Pavlov N, Wallbank JA, Hermans SM, Kingsbury JM, Pantos O, Lear G. 2025. Putative plastic-degrading communities within New Zealand’s geothermal environments. Total Environ Microbiol 1:100012.

15. Wilson CJN, Houghton BF, McWilliams MO, Lanphere MA, Weaver SD, Briggs RM. 1995. Volcanic and structural evolution of the Taupo Volcanic Zone, New Zealand: a review. J Volcanol Geotherm Res 68:1–28.

16. Power JF, Carere CR, Lee CK, Wakerley GLJ, Evans DW, Button M, White D, Climo MD, Hinze AM, Morgan XC, McDonald IR, Cary SC, Stott MB. 2018. Microbial biogeography of 925 geothermal springs in New Zealand. Nat Commun 9:2876.

17. Hetzer A, McDonald IR, Morgan HW. 2008. Venenivibrio stagnispumantis gen. nov., sp. nov., a thermophilic hydrogen-oxidizing bacterium isolated from Champagne Pool, Waiotapu, New Zealand. Int J Syst Evol Microbiol 58:398–403.

18. Greening C, Carere CR, Rushton-Green R, Harold LK, Hards K, Taylor MC, Morales SE, Stott MB, Cook GM. 2015. Persistence of the dominant soil phylum Acidobacteria by trace gas scavenging. Proc Natl Acad Sci U S A 112:10497–10502.

19. Anders H, Power JF, MacKenzie AD, Lagutin K, Vyssotski M, Hanssen E, Moreau JW, Stott MB. 2015. Limisphaera ngatamarikiensis gen. nov., sp. nov., a thermophilic, pink- pigmented coccus isolated from subaqueous mud of a geothermal hot spring. Int J Syst Evol Microbiol 65:1114–1121.

20. Leung J, Verwilligen V. 2023. TPMS Studio (version 0.8.4). University of Canterbury, Biomolecular Interaction Centre.

21. Dragone NB, Childress MK, Vanderburgh C, Hogg ID, Sancho L, Lee CK, Barrett JE, LeMonte J, Quandt CA, Fierer N. 2025. A comprehensive survey of microbial diversity across the Antarctic continent. Polar Biol 48:50.

22. Walters W, Hyde ER, Berg-Lyons D, Ackermann G, Humphrey G, Parada A, Gilbert JA, Jansson JK, Caporaso JG, Fuhrman JA, Apprill A, Knight R. 2016. Improved bacterial 16S rRNA gene (V4 and V4–5) and fungal internal transcribed spacer marker gene primers for microbial community surveys. mSystems 1:e00009–15.

23. Thompson LR, Sanders JG, McDonald D, Amir A, Ladau J, Locey KJ, Prill RJ, Tripathi A, Gibbons SM, Ackermann G, Navas-Molina JA, Janssen S, Kopylova E, Vázquez-Baeza Y, González A, Morton JT, Mirarab S, Zech Xu Z, Jiang L, Haroon MF, Kanbar J, Zhu Q, Jin Song S, Kosciolek T, Bokulich NA, Lefler J, Brislawn CJ, Humphrey G, Owens SM, Hampton-Marcell J, Berg-Lyons D, McKenzie V, Fierer N, Fuhrman JA, Clauset A, Stevens RL, Shade A, Pollard KS, Goodwin KD, Jansson JK, Gilbert JA, Knight R. 2017. A communal catalogue reveals Earth’s multiscale microbial diversity. Nature 551:457–463.

24. Dragone NB, Whittaker K, Lord OM, Burke EA, Dufel H, Hite E, Miller F, Page G, Slayback D, Fierer N. 2023. The early microbial colonizers of a short-lived volcanic island in the Kingdom of Tonga. mBio 14:e03313–22.

25. Dragone NB, Henley JB, Holland-Moritz H, Diaz M, Hogg ID, Lyons WB, Wall DH, Adams BJ, Fierer N. 2022. Elevational constraints on the composition and genomic attributes of microbial communities in Antarctic soils. mSystems 7:e01330–21.

26. Caporaso JG, Lauber CL, Walters WA, Berg-Lyons D, Huntley J, Fierer N, Owens SM, Betley J, Fraser L, Bauer M, Gormley N, Gilbert JA, Smith G, Knight R. 2012. Ultra-high-throughput microbial community analysis on the Illumina HiSeq and MiSeq platforms. ISME J 6:1621–1624.

27. Buchner D. 2022. PCR normalization and size selection with magnetic beads. https://www.protocols.io/view/pcr-normalization-and-size-selection-with-magnetic-ch4wt8xe

28. Callahan BJ, McMurdie PJ, Rosen MJ, Han AW, Johnson AJA, Holmes SP. 2016. DADA2: high-resolution sample inference from Illumina amplicon data. Nat Methods 13:581–583.

29. Wang Q, Garrity GM, Tiedje JM, Cole JR. 2007. Naïve Bayesian classifier for rapid assignment of rRNA sequences into the new bacterial taxonomy. Appl Environ Microbiol 73:5261–5267.

30. Yilmaz P, Parfrey LW, Yarza P, Gerken J, Pruesse E, Quast C, Schweer T, Peplies J, Ludwig W, Glöckner FO. 2014. The SILVA and “all-species living tree project (LTP)” taxonomic frameworks. Nucleic Acids Res 42:D643–D648.

31. Quast C, Pruesse E, Yilmaz P, Gerken J, Schweer T, Yarza P, Peplies J, Glöckner FO. 2013. The SILVA ribosomal RNA gene database project: improved data processing and web-based tools. Nucleic Acids Res 41:D590–D596.

32. R Core Team. 2023. R: a language and environment for statistical computing. R Foundation for Statistical Computing, Vienna, Austria.

33. PJ, Holmes S. 2013. phyloseq: an R package for reproducible interactive analysis and graphics of microbiome census data. PLoS One 8:e61217.

34. Zhang H, Perez-Garcia P, Dierkes RF, Applegate V, Schumacher J, Chibani CM, Sternagel S, Preuss L, Weigert S, Schmeisser C, Danso D, Pleiss J, Almeida A, Höcker B, Hallam SJ, Schmitz RA, Smits SHJ, Chow J, Streit WR. 2022. The Bacteroidetes Aequorivita sp. and Kaistella jeonii produce promiscuous esterases with PET-hydrolyzing activity. Front Microbiol 12:803896.

35. Khan S, Ali SA, Ali AS. 2023. Biodegradation of low-density polyethylene (LDPE) by the mesophilic fungus Penicillium citrinum isolated from soils of a plastic waste dump yard, Bhopal, India. Environ Technol 44:2300–2314.

36. Chien H-L, Tsai Y-T, Tseng W-S, Wu J-A, Kuo S-L, Chang S-L, Huang S-J, Liu C-T. 2022. Biodegradation of PBSA films by elite Aspergillus isolates and farmland soil. Polymers 14:1320.

37. Bolger AM, Lohse M, Usadel B. 2014. Trimmomatic: a flexible trimmer for Illumina sequence data. Bioinformatics 30:2114–2120.

38. Wick RR, Judd LM, Gorrie CL, Holt KE. 2017. Unicycler: resolving bacterial genome assemblies from short and long sequencing reads. PLoS Comput Biol 13:e1005595.

39. Parks DH, Imelfort M, Skennerton CT, Hugenholtz P, Tyson GW. 2015. CheckM: assessing the quality of microbial genomes recovered from isolates, single cells, and metagenomes. Genome Res 25:1043–1055.

40. Hyatt D, Chen G-L, LoCascio PF, Land ML, Larimer FW, Hauser LJ. 2010. Prodigal: prokaryotic gene recognition and translation initiation site identification. BMC Bioinformatics 11:119.

41. Chaumeil P-A, Mussig AJ, Hugenholtz P, Parks DH. 2020. GTDB-Tk: a toolkit to classify genomes with the Genome Taxonomy Database. Bioinformatics 36:1925–1927.

42. Price MN, Dehal PS, Arkin AP. 2009. FastTree: computing large minimum evolution trees with profiles instead of a distance matrix. Mol Biol Evol 26:1641–1650.

43. Parks DH, Chuvochina M, Rinke C, Mussig AJ, Chaumeil P-A, Hugenholtz P. 2022. GTDB: an ongoing census of bacterial and archaeal diversity through a phylogenetically consistent, rank normalized, and complete genome-based taxonomy. Nucleic Acids Res 50:D785–D794.

44. Gruber-Vodicka HR, Seah BKB, Pruesse E. 2020. phyloFlash: rapid small-subunit rRNA profiling and targeted assembly from metagenomes. mSystems 5:e00920–20.

45. Glöckner FO, Yilmaz P, Quast C, Gerken J, Beccati A, Ciuprina A, Bruns G, Yarza P, Peplies J, Westram R, Ludwig W. 2017. Twenty-five years of serving the community with ribosomal RNA gene reference databases and tools. J Biotechnol 261:169–176.

46. Letunic I, Bork P. 2024. Interactive Tree Of Life (iTOL) v6: recent updates to the phylogenetic tree display and annotation tool. Nucleic Acids Res 52:W78–W82.

47. Dragone NB, van Hamelsveld S, Nazmi AR, Stott MB, Hatley GA, Moloney K, Bohm K, Gutiérrez Ginés MJ, Weaver L. 2025. Examining the potential of plastic-fed black soldier fly larvae (Hermetia illucens) as “bioincubators” of plastic-degrading bacteria. J Appl Microbiol 136:lxaf085.

48. Gan Z, Zhang H. 2019. PMBD: a comprehensive plastics microbial biodegradation database. Database 2019:baz119.

49. Buchholz PCF, Feuerriegel G, Zhang H, Perez-Garcia P, Nover L-L, Chow J, Streit WR, Pleiss J. 2022. Plastics degradation by hydrolytic enzymes: the plastics-active enzymes database—PAZy. Proteins 90:1443–1456.

50. Galperin MY, Wolf YI, Makarova KS, Vera Alvarez R, Landsman D, Koonin EV. 2021. COG database update: focus on microbial diversity, model organisms, and widespread pathogens. Nucleic Acids Res 49:D274–D281.

51. Kanehisa M, Goto S. 2000. KEGG: Kyoto encyclopedia of genes and genomes. Nucleic Acids Res 28:27–30.

52. Ashburner M, Ball CA, Blake JA, Botstein D, Butler H, Cherry JM, Davis AP, Dolinski K, Dwight SS, Eppig JT, Harris MA, Hill DP, Issel-Tarver L, Kasarskis A, Lewis S, Matese JC, Richardson JE, Ringwald M, Rubin GM, Sherlock G. 2000. Gene ontology: tool for the unification of biology. Nat Genet 25:25–29.

53. The Gene Ontology Consortium, Aleksander SA, Balhoff J, Carbon S, Cherry JM, Drabkin HJ, Ebert D, Feuermann M, Gaudet P, Harris NL, Hill DP, Lee R, Mi H, Moxon S, Mungall CJ, Muruganugan A, Mushayahama T, Sternberg PW, Thomas PD, Van Auken K, Ramsey J, Siegele DA, Chisholm RL, Fey P, Aspromonte MC, Nugnes MV, Quaglia F, Tosatto S, Giglio M, Nadendla S, Antonazzo G, Attrill H, dos Santos G, Marygold S, Strelets V, Tabone CJ, Thurmond J, Zhou P, Ahmed SH, Asanitthong P, Luna Buitrago D, Erdol MN, Gage MC, Ali Kadhum M, Li KYC, Long M, Michalak A, Pesala A, Pritazahra A, Saverimuttu SCC, Su R, Thurlow KE, Lovering RC, Logie C, Oliferenko S, Blake J, Christie K, Corbani L, Dolan ME, Ni L, Sitnikov D, Smith C, Cuzick A, Seager J, Cooper L, Elser J, Jaiswal P, Gupta P, Naithani S, Lera-Ramirez M, Rutherford K, Wood V, De Pons JL, Dwinell MR, Hayman GT, Kaldunski ML, Kwitek AE, Laulederkind SJF, Tutaj MA, Vedi M, Wang S-J, D’Eustachio P, Aimo L, Axelsen K, Bridge A, Hyka-Nouspikel N, Morgat A, Engel SR, Karra K, Miyasato SR, Nash RS, Skrzypek MS, Weng S, Wong ED, Bakker E, Berardini TZ, Reiser L, Auchincloss A, Argoud-Puy G, Blatter M-C, Boutet E, Breuza L, Casals-Casas C, Coudert E, Estreicher A, Famiglietti ML, Gos A, Gruaz-Gumowski N, Hulo C, Jungo F, Le Mercier P, Lieberherr D, Masson P, Pedruzzi I, Pourcel L, Poux S, Rivoire C, Sundaram S, Bateman A, Bowler-Barnett E, Bye-A-Jee H, Denny P, Ignatchenko A, Ishtiaq R, Lock A, Lussi Y, Magrane M, Martin MJ, Orchard S, Raposo P, Speretta E, Tyagi N, Warner K, Zaru R, Diehl AD, Chan J, Diamantakis S, Raciti D, Zarowiecki M, Fisher M, James-Zorn C, Ponferrada V, Zorn A, Ramachandran S, Ruzicka L, Westerfield M. 2023. The Gene Ontology knowledgebase in 2023. Genetics 224:iyad031.

54. The UniProt Consortium. 2019. UniProt: a worldwide hub of protein knowledge. Nucleic Acids Res 47:D506–D515.

55. Rognes T, Flouri T, Nichols B, Quince C, Mahé F. 2016. VSEARCH: a versatile open source tool for metagenomics. PeerJ 4:e2584.

56. Wood SN. 2025. Generalized additive models. Annu Rev Stat Appl 12:497–526.

57. QGIS Development Team. 2025. QGIS geographic information system (version 3.44). Open Source Geospatial Foundation.

58. CARTO. 2026. CARTO Positron raster basemap tiles. CARTO.

59. Salter SJ, Cox MJ, Turek EM, Calus ST, Cookson WO, Moffatt MF, Turner P, Parkhill J, Loman NJ, Walker AW. 2014. Reagent and laboratory contamination can critically impact sequence-based microbiome analyses. BMC Biol 12:87.

60. McAdam B, Fournet MB, McDonald P, Mojicevic M. 2020. Production of polyhydroxybutyrate (PHB) and factors impacting its chemical and mechanical characteristics. Polymers 12:2908.

61. Raza ZA, Abid S, Banat IM. 2018. Polyhydroxyalkanoates: characteristics, production, recent developments, and applications. Int Biodeterior Biodegrad 126:45–56.

62. Lemoigne M. 1926. Produit de deshydratation et de polymerisation de l’acide β-oxybutyrique. Bull Soc Chim Biol 8:770–782.

63. Bourque D, Ouellette B, André G, Groleau D. 1992. Production of poly-β-hydroxybutyrate from methanol: characterization of a new isolate of Methylobacterium extorquens. Appl Microbiol Biotechnol 37:7–12.

64. Ochsner AM, Sonntag F, Buchhaupt M, Schrader J, Vorholt JA. 2015. Methylobacterium extorquens: methylotrophy and biotechnological applications. Appl Microbiol Biotechnol 99:517–534.

65. Jendrossek D. 2009. Polyhydroxyalkanoate granules are complex subcellular organelles (carbonosomes). J Bacteriol 191:3195–3202.

66. Müller-Santos M, Koskimäki JJ, Alves LPS, de Souza EM, Jendrossek D, Pirttilä AM. 2021. The protective role of PHB and its degradation products against stress situations in bacteria. FEMS Microbiol Rev 45:fuaa058.

67. Dragone NB, Hoffert M, Strickland MS, Fierer N. 2024. Taxonomic and genomic attributes of oligotrophic soil bacteria. ISME Commun 4:ycae081.

68. Wang JL, Dragone NB, Avard G, Hynek BM. 2022. Microbial survival in an extreme Martian analog ecosystem: Poás Volcano, Costa Rica. Front Astron Space Sci 9:817900.

69. Bao Q, Zhang Z, Luo H, Tao X. 2023. Evaluating and modeling the degradation of PLA/PHB fabrics in marine water. Polymers 15:82.

70. Lyshtva P, Voronova V, Barbir J, Filho WL, Kröger SD, Witt G, Miksch L, Saborowski R, Gutow L, Frank C, Emmerstorfer-Augustin A, Agustin-Salazar S, Cerruti P, Santagata G, Stagnaro P, D’Arrigo C, Vignolo M, Krång A-S, Strömberg E, Lehtinen L Annunen V. 2024. Degradation of a poly(3-hydroxybutyrate-co-3-hydroxyvalerate) (PHBV) compound in different environments. Heliyon 10:e24770.

71. Benavides Fernández CD, Guzmán Castillo MP, Quijano Pérez SA, Carvajal Rodríguez LV. 2022. Microbial degradation of polyethylene terephthalate: a systematic review. SN Appl Sci 4:263.

72. Marqués-Calvo MS, Cerdà-Cuéllar M, Kint DPR, Bou JJ, Muñoz-Guerra S. 2006. Enzymatic and microbial biodegradability of poly(ethylene terephthalate) copolymers containing nitrated units. Polym Degrad Stab 91:663–671.

73. Mohanan N, Montazer Z, Sharma PK, Levin DB. 2020. Microbial and enzymatic degradation of synthetic plastics. Front Microbiol 11:580709.

74. Kawai F, Oda M, Tamashiro T, Waku T, Tanaka N, Yamamoto M, Mizushima H, Miyakawa T, Tanokura M. 2014. A novel Ca2+-activated, thermostabilized polyesterase capable of hydrolyzing polyethylene terephthalate from Saccharomonospora viridis AHK190. Appl Microbiol Biotechnol 98:10053–10064.

75. Rosli NA, Karamanlioglu M, Kargarzadeh H, Ahmad I. 2021. Comprehensive exploration of natural degradation of poly(lactic acid) blends in various degradation media: a review. Int J Biol Macromol 187:732–741.

76. Krasowska K, Heimowska A. 2023. Degradability of polylactide in natural aqueous environments. Water 15:198.

77. Itävaara M, Karjomaa S, Selin J-F. 2002. Biodegradation of polylactide in aerobic and anaerobic thermophilic conditions. Chemosphere 46:879–885.

78. Hahladakis JN, Velis CA, Weber R, Iacovidou E, Purnell P. 2018. An overview of chemical additives present in plastics: migration, release, fate, and environmental impact during their use, disposal, and recycling. J Hazard Mater 344:179–199.

79. Lear G, Maday SDM, Gambarini V, Northcott G, Abbel R, Kingsbury JM, Weaver L, Wallbank JA, Pantos O. 2022. Microbial abilities to degrade global environmental plastic polymer waste are overstated. Environ Res Lett 17:043002.

80. Brebu M. 2020. Environmental degradation of plastic composites with natural fillers—a review. Polymers 12:166.

81. Bunbury F, Rivas C, Calatrava V, Malkovskiy AV, Joubert L-M, Parvate AD, Evans JE, Grossman AR, Bhaya D. 2025. Cyanobacteria and Chloroflexota cooperate to structure light-responsive biofilms. Proc Natl Acad Sci U S A 122:e2423574122.

82. Kaul A, Böllmann A, Thema M, Kalb L, Stöckl R, Huber H, Sterner M, Bellack A. 2022. Combining a robust thermophilic methanogen and packing material with high liquid hold-up to optimize biological methanation in trickle-bed reactors. Bioresour Technol 345:126524.

83. Anda D, Makk J, Krett G, Jurecska L, Márialigeti K, Mádl-Szőnyi J, Borsodi AK. 2015. Thermophilic prokaryotic communities inhabiting the biofilm and well water of a thermal karst system located in Budapest (Hungary). Extremophiles 19:787–797.

84. Jabeen I, Islam S, Hassan AKMI, Tasnim Z, Shuvo SR. 2023. A brief insight into Citrobacter species—a growing threat to public health. Front Antibiot 2:1276982.

85. Di Gregorio L, Tandoi V, Congestri R, Rossetti S, Di Pippo F. 2017. Unravelling the core microbiome of biofilms in cooling tower systems. Biofouling 33:793–806.

86. De Filippis F, Bonelli M, Bruno D, Sequino G, Montali A, Reguzzoni M, Pasolli E, Savy D, Cangemi S, Cozzolino V, Tettamanti G, Ercolini D, Casartelli M, Caccia S. 2023. Plastics shape the black soldier fly larvae gut microbiome and select for biodegrading functions. Microbiome 11:205.

87. Amaral-Zettler LA, Zettler ER, Mincer TJ. 2020. Ecology of the plastisphere. Nat Rev Microbiol 18:139–151.

88. Lear G, Kingsbury JM, Franchini S, Gambarini V, Maday SDM, Wallbank JA, Weaver L, Pantos O. 2021. Plastics and the microbiome: impacts and solutions. Environ Microbiome 16:2.

89. Lu J, Takahashi A, Ueda S. 2014. 3-hydroxybutyrate oligomer hydrolase and 3-hydroxybutyrate dehydrogenase participate in intracellular polyhydroxybutyrate and polyhydroxyvalerate degradation in Paracoccus denitrificans. Appl Environ Microbiol 80:986–993.

90. Zhou W, Bergsma S, Colpa DI, Euverink G-JW, Krooneman J. 2023. Polyhydroxyalkanoates (PHAs) synthesis and degradation by microbes and applications towards a circular economy. J Environ Manag 341:118033.

91. Brigham CJ, Reimer EN, Rha C, Sinskey AJ. 2012. Examination of PHB depolymerases in Ralstonia eutropha: further elucidation of the roles of enzymes in PHB homeostasis. AMB Express 2:26.

92. Jang JW, Hwang IY, Lee OK, Lee EY. 2025. Production of polyhydroxybutyrate with high cell density cultivation using the thermophile Caldimonas thermodepolymerans. Bioresour Technol 419:132073.

93. An J, Ha B, Lee SK. 2023. Production of polyhydroxyalkanoates by the thermophile Cupriavidus cauae PHS1. Bioresour Technol 371:128627.

94. Ariza-Tarazona MC, Villarreal-Chiu JF, Hernández-López JM, Rivera De la Rosa J, Barbieri V, Siligardi C, Cedillo-González EI. 2020. Microplastic pollution reduction by a carbon- and nitrogen-doped TiO2: effect of pH and temperature in the photocatalytic degradation process. J Hazard Mater 395:122632.

