## Supplemental Information for "Isolation of thermophilic plastic-degrading bacteria from hot springs of Aotearoa-New Zealand"

*Applied and Environmental Microbiology*

…

**Contents of this file**

Supplementary Methods

Tables S1 – S4

Figures S1 to S8

Captions for Dataset S1 – S4

Supplemental References 1 - 2

**Additional Supporting Information (files uploaded separately)**

Datasets S1 – S4

**Summary**

The supporting information in this file contains supplemental methods, tables, figures, and captions for attached data sets referenced in the main paper. All processing steps used to generate these data are described in the methods. More specifically this SI provide more specific details about the media used to isolate our organisms, the springs used for the incubation studies and their geochemical properties, the results of 16S rRNA gene amplicon sequencing, the isolation and characterization of our plastic degrading organisms, and the predictive modeling of the thermophilic plastic degrading organisms.

**Supplementary Methods: Media Recipe (Modified from DSMZ 150a: *Acidothiobacillus caldus* medium)**

**Media Recipe**

| **Component** | **Amount per Liter** |
| --- | --- |
| (NH_4_)_2_SO_4_ | 1.0 g |
| K_2_HPO_4_ x 3 H_2_O | 0.50 g |
| MgSO_4_ x 7 H_2_O | 0.50 g |
| KCl | 0.10 g |
| Ca(NO_3_)_2_ x 4 H_2_O | 0.02g |
| Noble Agar | 2 g |
| Chelated Iron Solution | 3ml |
| Methanotroph trace element solution | 1ml |
| Methanogen trace element solution | 1ml |
| Distilled H_2_O | 995 ml |

**Chelated iron Solution (DSMZ 748 CNES Medium)**

| **Component** | **Amount per Liter** |
| --- | --- |
| FeCl_2_ x 4 H_2_O | 1.0g |
| EDTA | 2.0g |
| HCl (concentrated) | 3ml |
| Distilled H_2_O | 900ml |

**Methanotroph trace element solution (From DSMZ 632 NMS medium):**

| **Component** | **Amount per liter** |
| --- | --- |
| Na_2_-EDTA | 500.0 mg |
| FeSO_4_ x 7 H_2_O | 200.0 mg |
| ZnSO_4_ x 7 H_2_O | 10.0 mg |
| MnCl_2_ x 4 H_2_O | 3.0 mg |
| H_3_BO_3_ | 30.0 mg |
| CoCl_2_ x 6 H_2_O | 20.0 mg |
| CaCl_2_ x 2 H_2_O | 1.0 mg |
| NiCl_2_ x 6 H_2_O | 2.0 mg |
| Na_2_MoO_4_ x 2 H_2_O | 3.0 mg |
| Distilled H_2_O | 1000ml |

**Methanogen trace element solution SL-10 (From DSMZ 119 methanobacterium medium)**

| **Component:** | **Amount per liter** |
| --- | --- |
| HCl (25%) | 10.0 ml |
| FeCl_2_ x 4 H_2_O | 1.5 g |
| ZnCl_2_ | 70mg |
| MnCl_2_ x 4 H_2_O | 100 mg |
| H_3_BO_3_ | 6.0 mg |
| CoCl_3_ x 6 H_2_O | 190 mg |
| CuCl_2_ x 2 H_2_O | 2.0 |
| NiCl_2_ x 6 H_2_O | 24.0 mg |
| Na_2_MoO_4_ x 2 H_2_O | 36.0 mg |
| Distilled H_2_O | 990 ml |

**Instructions:**

Dissolve ingredients, minus trace element solution and agar, in distilled H_2_O so the total volume is 1000ml. Add just enough noble agar to ensure plates will solidify (2% w/v of media). To ensure the noble agar is fully dissolved, bring solution to a boil while stirring constantly to avoid clumping. Adjust pH as required then autoclave media to sterilize. Once cooled, add filter sterilized chelated iron solution (3 mL/L), methanotroph trace element solution (3 mL/L), and methanogen trace element solutions (1 mL/L) was added to each media aliquot

**Plastic overlay amendment**

| **Component** | **Amount** |
| --- | --- |
| Powdered plastic polymer | 10g |

**Instructions:**

Add powdered plastic to 250ml aliquot of media (prepared as above) before autoclaving. After sterilization and addition of trace element solutions, let cool to 55^o^C. Place back on stir plate to ensure mixing of plastic in media. Pipette thin layer (~10 ml for a 58 cm^2^ Petri dishes petri dish) of media on top of pre-poured and cooled plates. Let solidify.

**Table S1:** Top 10 most abundant taxa identified through the 16S rRNA gene sequencing in: **A)** All samples. **B)** Springs with pH >7.0. **C)** Springs with pH <7.0)

**A.**

Bacteria; Bacillota; Thermoanaerobacteria; Thermoanaerobacterales; Family III; Thermoanaerobacterium; ASV_1

Archaea; Methanobacteriota; Methanobacteria; Methanobacteriales; Methanothermobacteriaceae; Methanothermobacter; ASV_2

Archaea; Methanobacteriota; Methanobacteria; Methanobacteriales; Methanothermobacteriaceae; Methanothermobacter; ASV_3

Bacteria; Spirochaetota; Spirochaetia; Spirochaetales; Spirochaetaceae; Rectinema; ASV_6

Bacteria; Acetothermia; Acetothermiia; KB1 group; NA; NA; ASV_5

Bacteria; Bacillota; Negativicutes; Veillonellales-Selenomonadales; NA; NA; ASV_7

Bacteria; Nitrospirota; Thermodesulfovibrionia; Thermodesulfovibrionales; Thermodesulfovibrionaceae; Thermodesulfovibrio; ASV_8

Archaea; Halobacteriota; Methanosarcinia; Methanosarcinales; Methanosaetaceae; Methanothrix; ASV_9

Bacteria; Spirochaetota; Spirochaetia; Spirochaetales; Spirochaetaceae; Rectinema; ASV_10

Bacteria; Bacillota; Syntrophomonadia; Syntrophomonadales; Syntrophomonadaceae; Syntrophothermus; ASV_11

Bacteria; Bacillota; Thermoanaerobacteria; Thermoanaerobacterales; Family III; Thermoanaerobacterium; ASV_1

**B.**

Archaea; Thermoplasmatota; Thermoplasmata; A10; NA; NA;ASV_14

Archaea; Thermoproteota; Thermoproteia; Fervidicoccales; Fervidicoccaceae; Fervidicoccus; ASV_71

Archaea; Thermoplasmatota; Thermoplasmata; A10; NA; NA; ASV_82

Archaea; Thermoproteota; Bathyarchaeia; NA; NA; NA; ASV_75

Archaea; Thermoproteota; Thermoproteia; Thermoproteales; Thermoproteaceae; Pyrobaculum; ASV_59

Bacteria; Thermodesulfobacteriota; Thermodesulfobacteria; Thermodesulfobacteriales; Thermodesulfobacteriaceae; Caldimicrobium; ASV_115

Archaea; Thermoproteota; Bathyarchaeia; NA; NA; NA; ASV_141

Bacteria; Aquificota; Aquificia; Hydrogenothermales; Hydrogenothermaceae; Venenivibrio; ASV_101

Archaea; Korarchaeota; Korarchaeia; Korarchaeales; Korarchaeaceae; Candidatus Korarchaeum; ASV_165

Archaea; Thermoproteota; Thermoproteia; Thermoproteales; Thermofilaceae; Thermofilum; ASV_102

**C.**

Bacteria; Bacillota; Thermoanaerobacteria; Thermoanaerobacterales; Family III; Thermoanaerobacterium; ASV_1

Archaea; Methanobacteriota; Methanobacteria; Methanobacteriales; Methanothermobacteriaceae; Methanothermobacter; ASV_2

Archaea; Methanobacteriota; Methanobacteria; Methanobacteriales; Methanothermobacteriaceae; Methanothermobacter; ASV_3

Bacteria; Spirochaetota; Spirochaetia; Spirochaetales; Spirochaetaceae; Rectinema; ASV_6

Bacteria; Acetothermia; Acetothermiia; KB1 group; NA; NA; ASV_5

Bacteria; Bacillota; Negativicutes; Veillonellales-Selenomonadales; NA; NA; ASV_7

Bacteria; Nitrospirota; Thermodesulfovibrionia; Thermodesulfovibrionales; Thermodesulfovibrionaceae; Thermodesulfovibrio; ASV_8

Archaea; Halobacteriota; Methanosarcinia; Methanosarcinales; Methanosaetaceae; Methanothrix; ASV_9

Bacteria; Spirochaetota; Spirochaetia; Spirochaetales; Spirochaetaceae; Rectinema; ASV_10

Bacteria; Bacillota; Syntrophomonadia; Syntrophomonadales; Syntrophomonadaceae; Syntrophothermus; ASV_11

Bacteria; Bacillota; Thermoanaerobacteria; Thermoanaerobacterales; Family III; Thermoanaerobacterium; ASV_1

**Table S2:** Genera identified through indicator taxa analysis who have previously been reported to degrade plastic. Identification of plastic degrading organisms is based on PlasticDB (1) For more details, see methods.

| **ASV** | **Genus** | **Plastic Degraded** |
| --- | --- | --- |
| ASV_181 | Citrobacter | LDPE, PS |
| ASV_243 | Lysinibacillus | PE,PCL |
| ASV_456 | Corynebacterium | PU |
| ASV_407 | Pseudomonas | PHO,PVA,PU,P3HP,P4HB,PEA,PES,PHB,O-PVA,PCL,PHA,PHBV,PHPV,PBSA,PE,PBAT,  PET,PLA,PS,PVC Blend,HDPE,LDPE,PU Blend,  P3HO,PP,PVC,PBS,P(3HB-co-3MP),PHC,  P(3HB-co-4HB),PHA Blend,PHN,PS Blend,PEG |

**Table S3:** Results of the random forest models. Variables that were considered predictors of thermophilic plastic degrader abundance (*) if they increased the mean standard error percent (MSE%) by >5% with a pval of <0.05.

| **Variable** | **%IncMSE** | **%IncMSE.pval** |
| --- | --- | --- |
| pH* | 6.9711721 | 0.00990099 |
| Al | 4.7658586 | 0.03960396 |
| Sr | 4.3501282 | 0.16831683 |
| sulfate | 4.229214 | 0.03960396 |
| Cs | 3.7530195 | 0.34653465 |
| Rb | 3.6641577 | 0.30693069 |
| Li | 3.4748158 | 0.40594059 |
| Ca | 3.4368019 | 0.45544554 |
| turbidity | 3.3829956 | 0.28712871 |
| Si | 3.2466249 | 0.44554455 |
| B | 3.2206655 | 0.46534653 |
| As | 3.0555029 | 0.63366337 |
| K | 3.0469293 | 0.69306931 |
| Mn | 2.9866116 | 0.65346535 |
| chloride | 2.8542575 | 0.67326733 |
| initialTemp | 2.7973862 | 0.17821782 |
| conductivity | 2.7433546 | 0.82178218 |
| nitrate | 2.6066771 | 0.52475248 |
| Br | 2.5924291 | 0.83168317 |
| phosphate | 2.5298224 | 0.30693069 |
| Cu | 2.2430548 | 0.40594059 |
| Na | 2.1338545 | 0.96039604 |
| Zn | 1.4602463 | 0.83168317 |
| Mg | 1.1752808 | 0.99009901 |
| redox | 1.162358 | 0.95049505 |
| ammonium | 1.0576878 | 0.86138614 |
| Ba | 0.3590875 | 0.78217822 |
| dO | -0.7339969 | 0.97029703 |
| nitrite | -1.7433441 | 0.99009901 |

**Table S4:** Diagnostic metrics for our generalized additive models (GAMs) that were used to predict under which condition abundance of thermophilic plastic degraders was maximized. Metrics were calculated with gam.check function of r package ‘mgcv’ (see methods for more details).

| **Variable** | **k’** | **edf** | **kindex** | **pval** |
| --- | --- | --- | --- | --- |
| pH | 4.0 | 3.07 | 0.97 | 0.17 |
| Al | 4.0 | 2.86 | 1.03 | 0.86 |

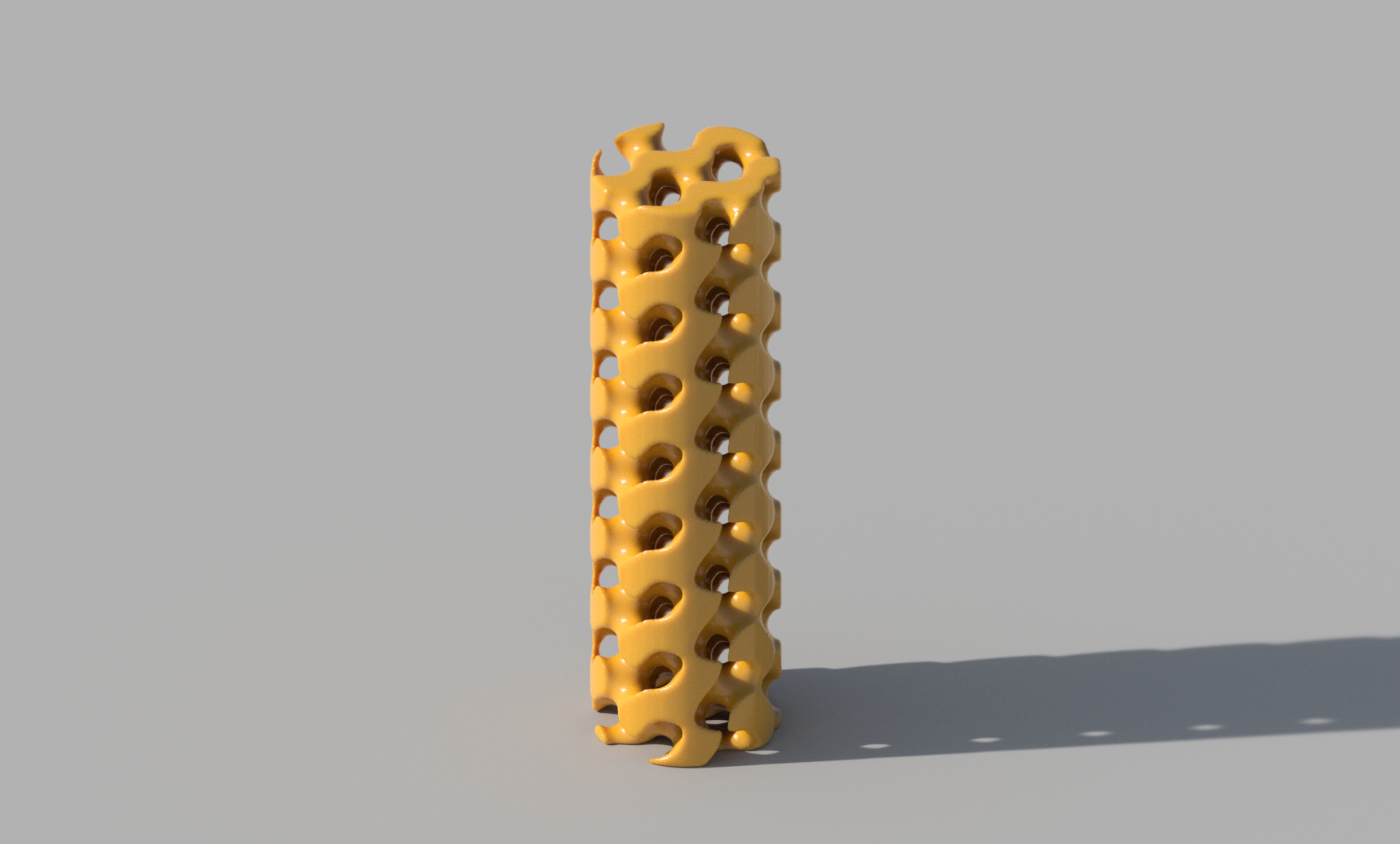

**Figure S1:** Digital model of the final TPMS gyroid structure used for this study.

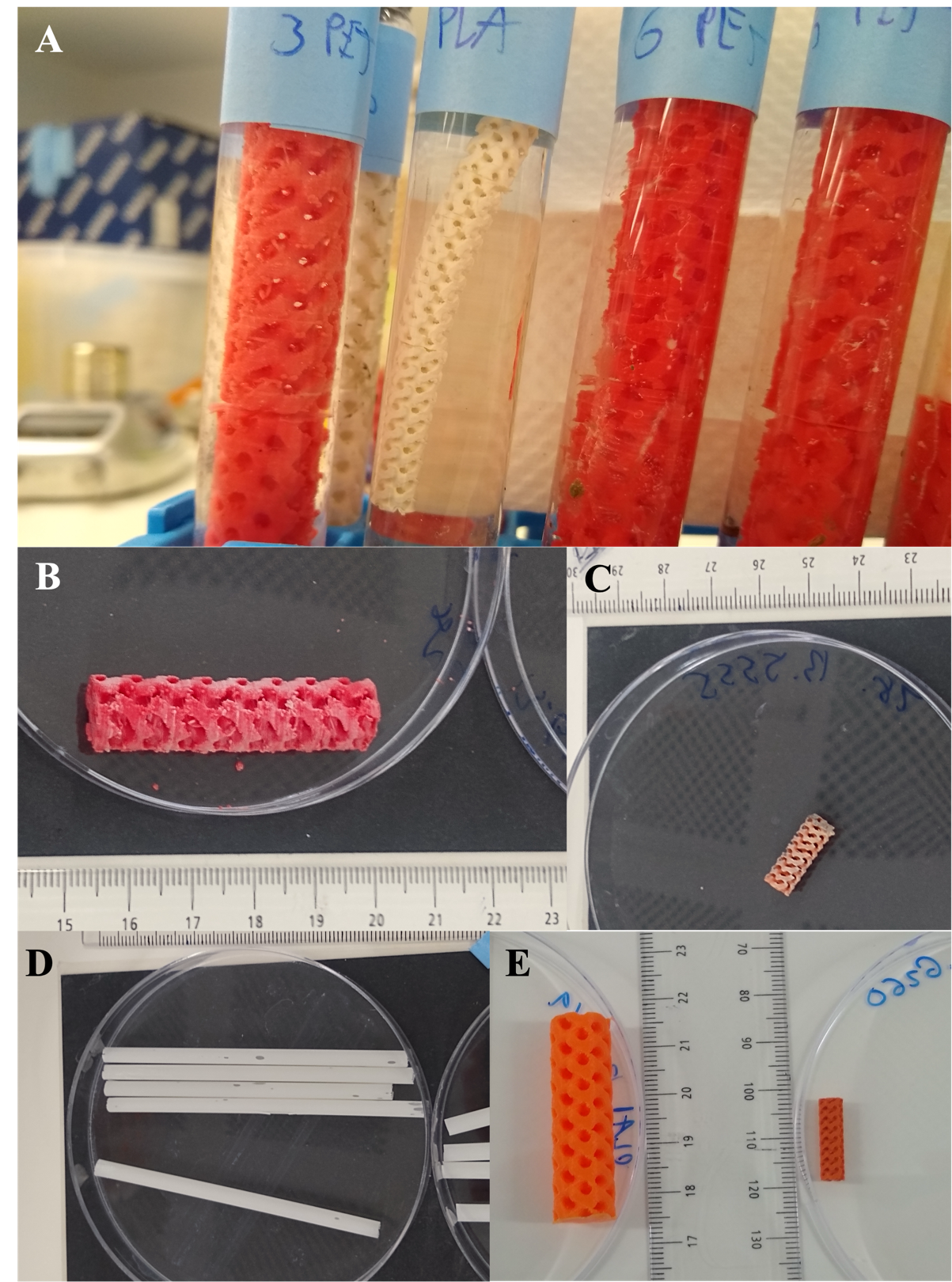

**Figure S2: Additional photos of substrate post-incubation. A)** Tubes removed from springs after three months. PLA tube in the center displays contraction and loss of coloring experienced at high temperatures. **B)** PET gyroid and **C)** PLA gyroid displaying color loss after incubation. **D)** Additional images of PLA straw degradation (See Figure 3 for another image). **E)** An unincubated gyroid (left) and post-incubation gyroid displaying contraction of the polymer (Right).

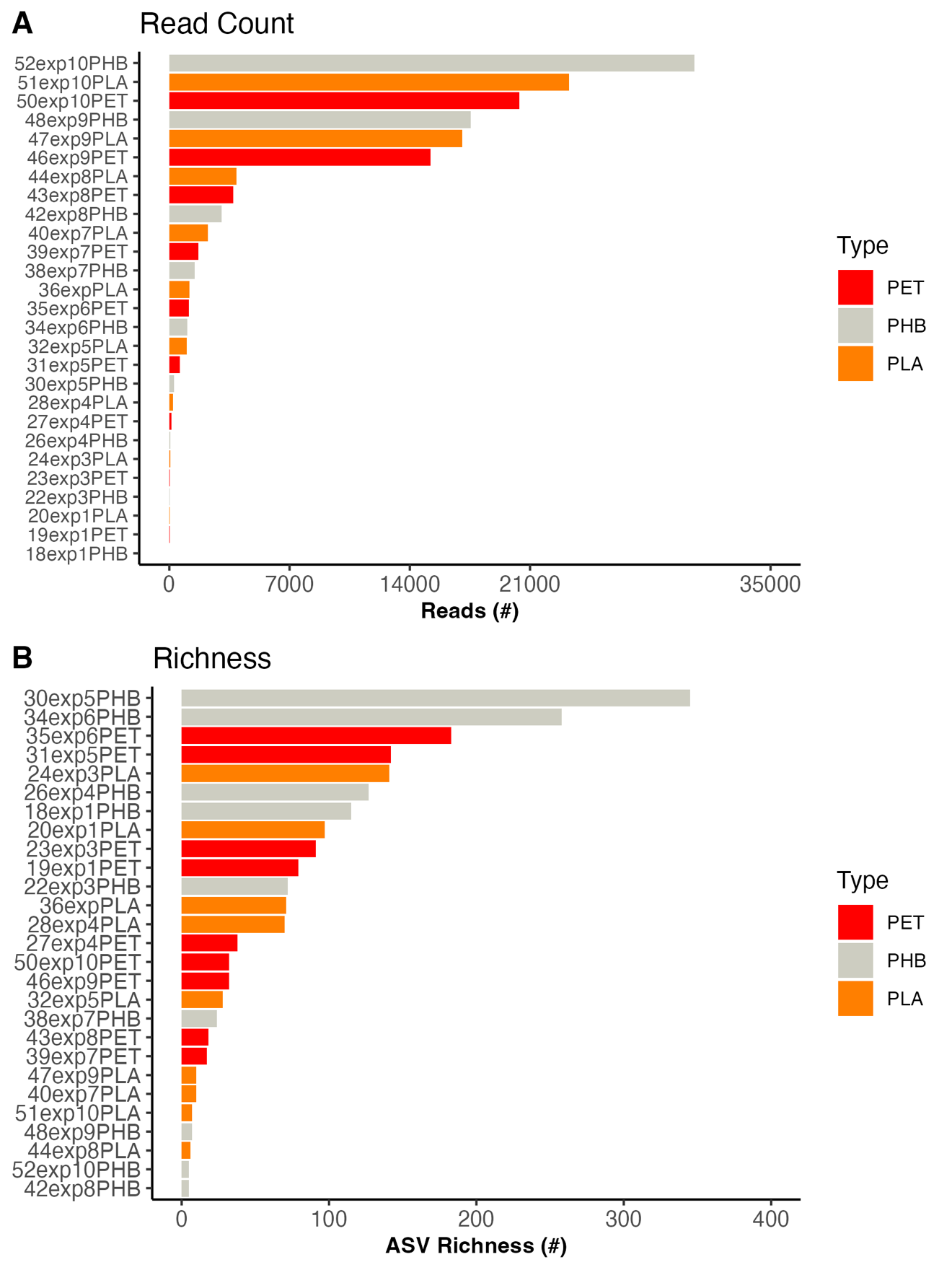

**Figure S3: A)** The number of reads recovered from the 27-incubation chamber through the 16S rRNA gene sequencing effort. **B)** The prokaryotic richness (number of distinct prokaryotic ASVs per sample) of the 27 incubation chambers. For both plots, samples are ordered from maximum read depth/richness to minimum and are colored by the plastic polymer contained.

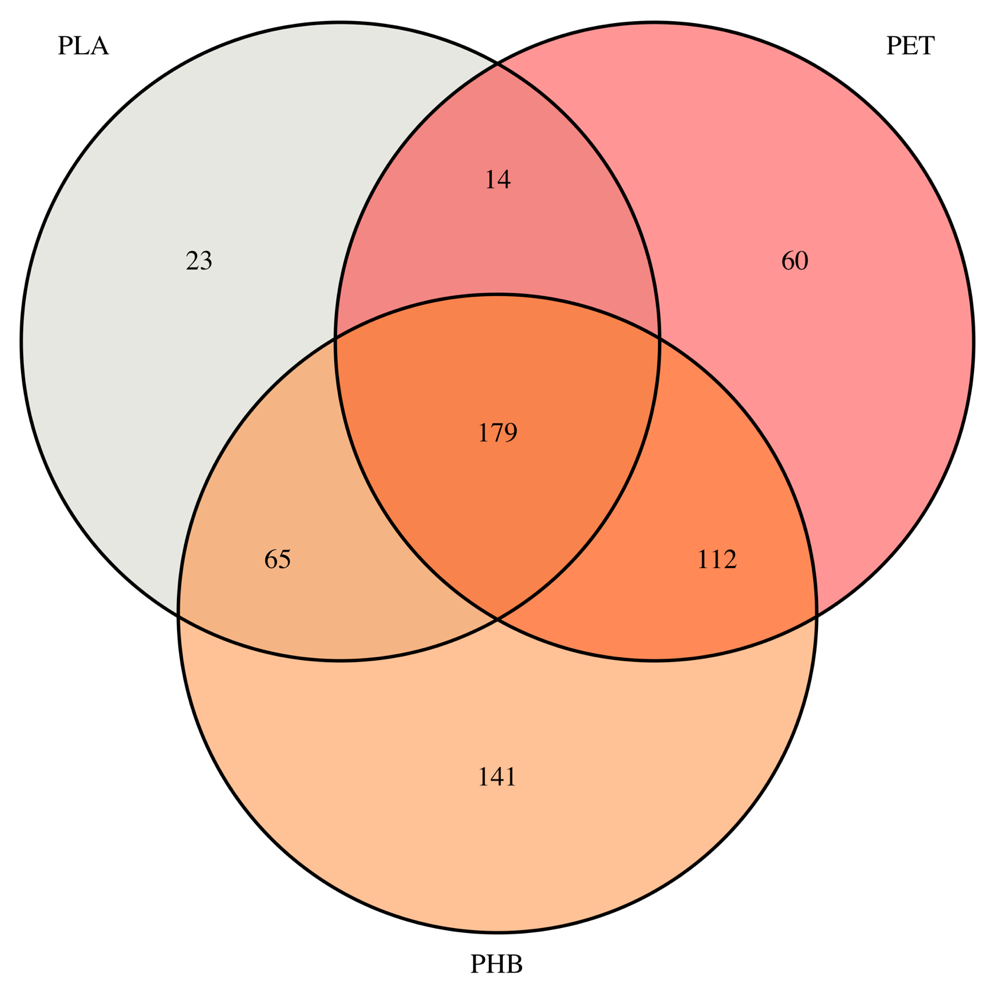

**Figure S4:** Shared prokaryotic taxa across the three polymer types. Venn diagram was created with the ASV list compiled from the 16S rRNA gene sequencing effort of the PHB gyroids (n = 9), PET (n = 9), PHA (n = 9). Venn diagram sets are colored based on the polymer type.

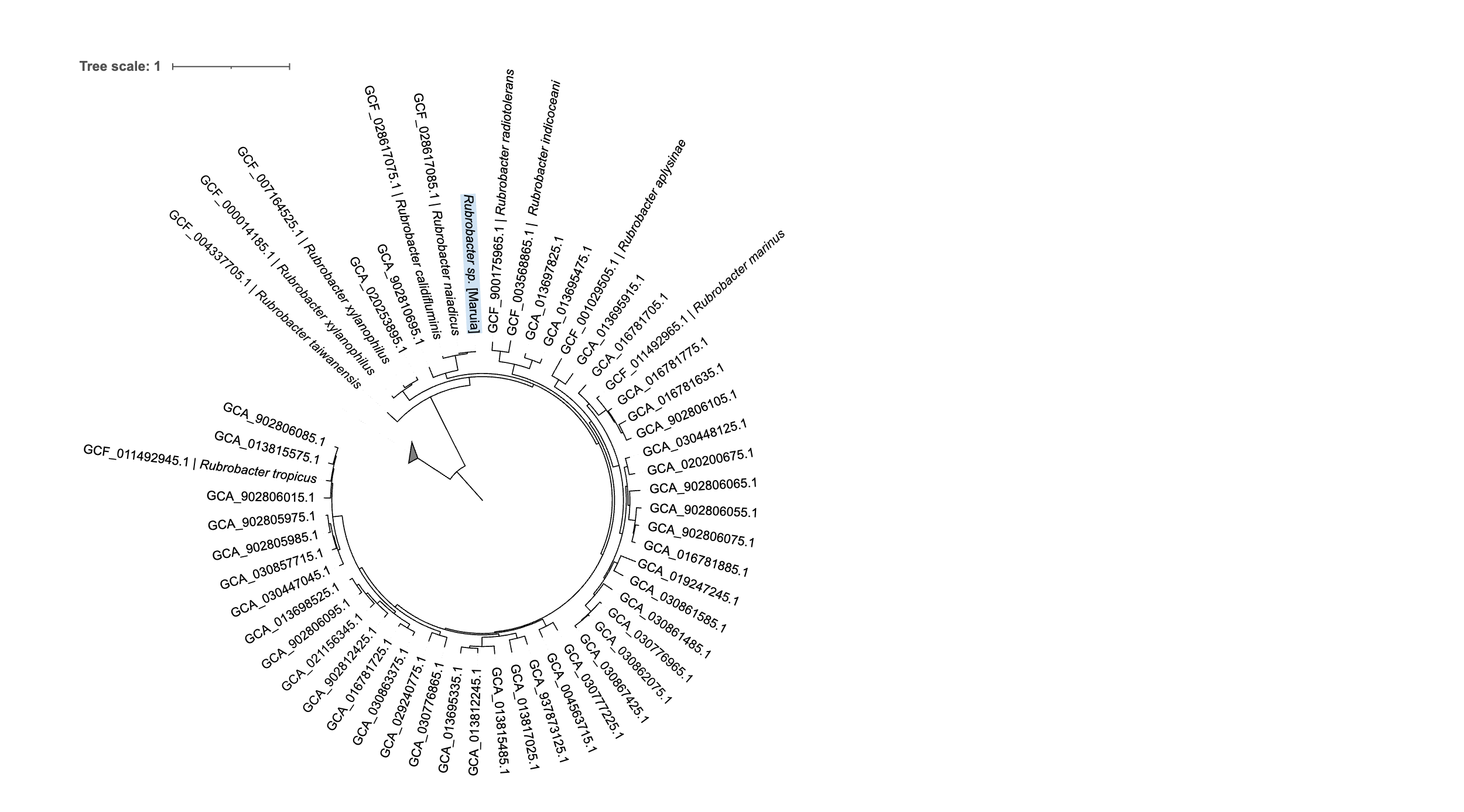

**Figure S5:** Phylogenetic tree placing our isolated *Rubrobacter sp.* [Maruia] within all GTDB reference sequences within the family Rubrobacteriaceae (53 genomes). The outgroup used to create the tree was the gtdb taxon group “g__Acidovorax.”. GTDB accession for all genomes is listed with our genome highlighted. If a genome is associated with a named type strain, the species is also listed.

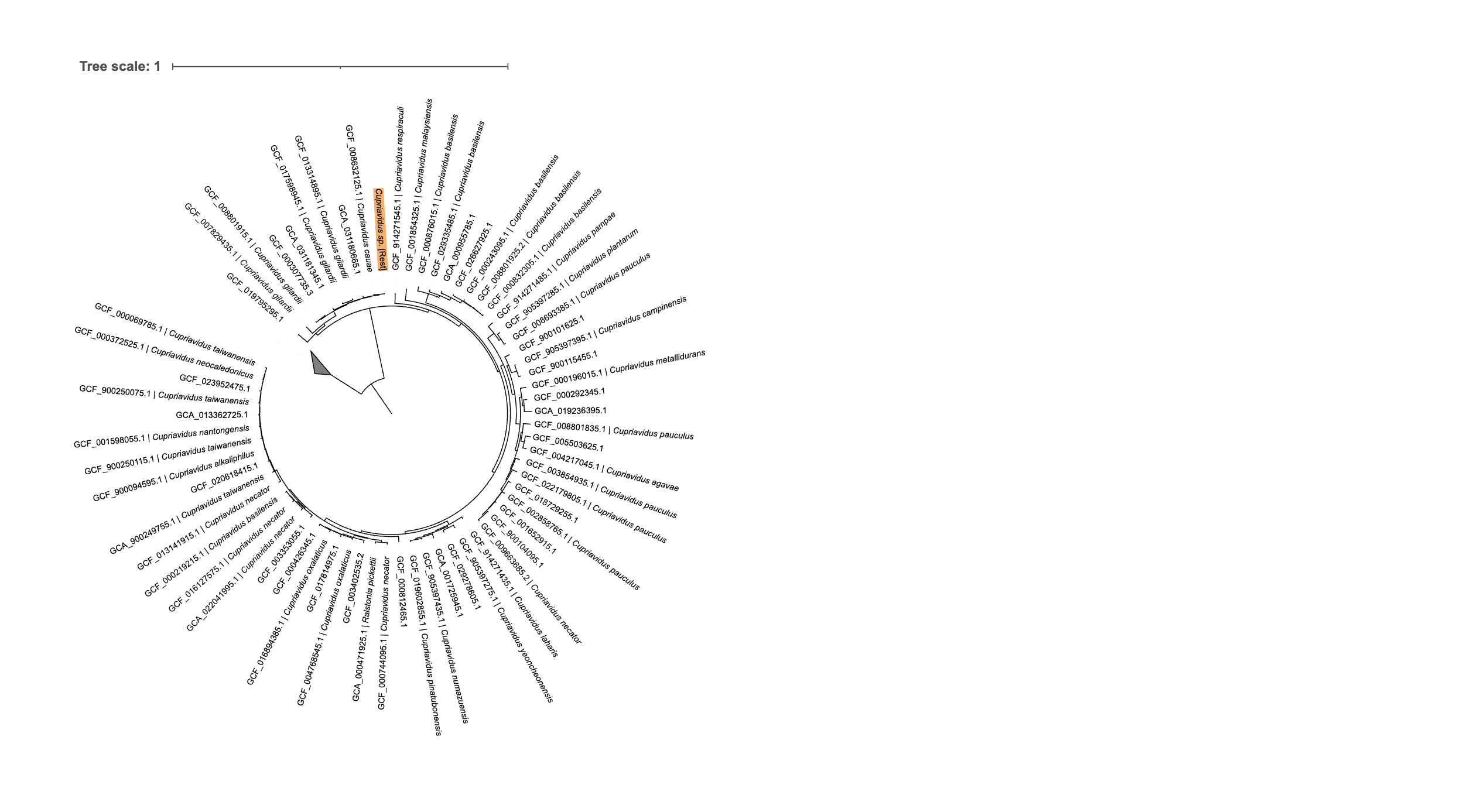

**Figure S6:** Phylogenetic tree placing our isolated *Cupriavidus sp* [Rest] within all GTDB reference sequences within the genus *Cupriavidus* (67 genomes). The outgroup used to create the tree was the gtdb taxon group “g__Acidovorax.” GTDB accession for all reference genomes is listed with our genome highlighted. If a genome is associated with a named type strain, the species is also listed.

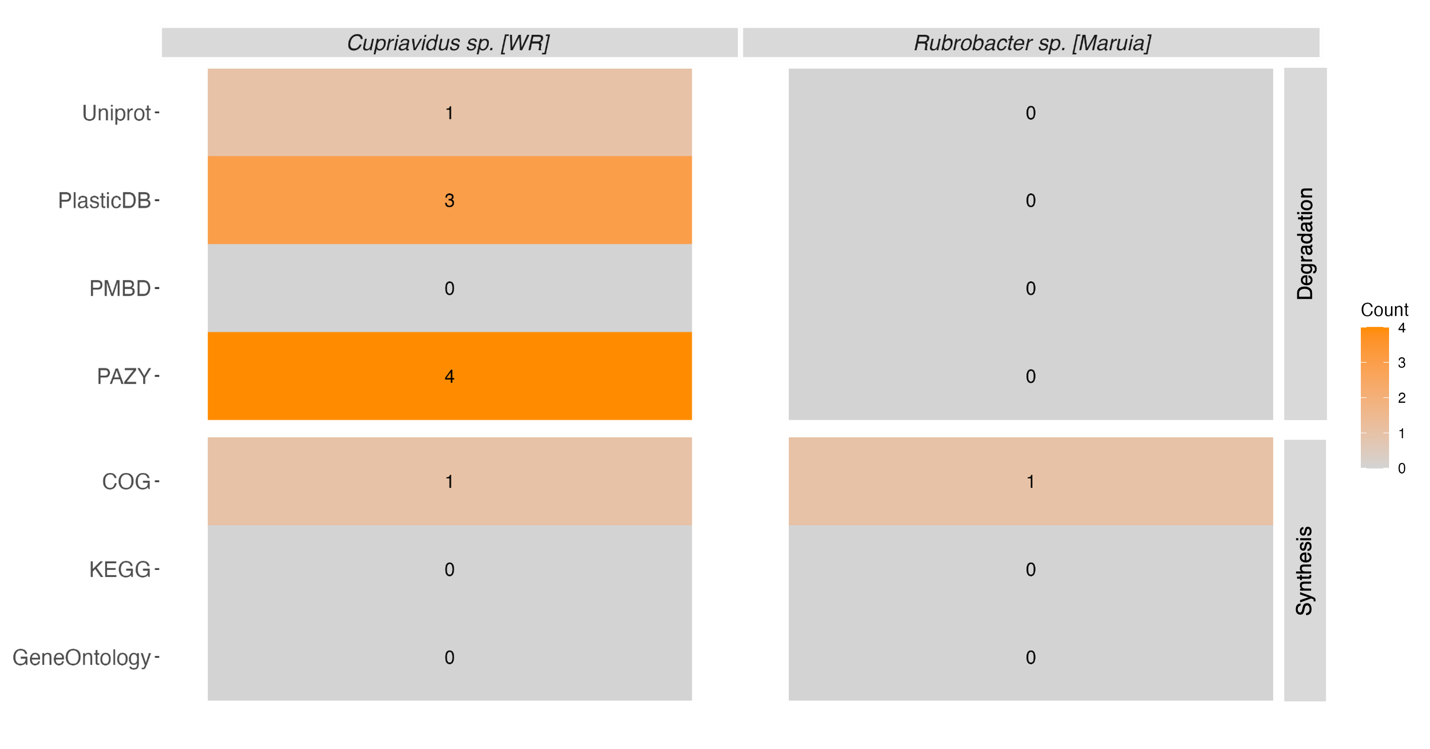

**Figure S7:** Heatmap displaying the number of hits to reference protein sequences related to PHA degradation and synthesis identified within our two isolate genomes. Hits are grouped based on the database that the reference sequences came from. For more information about the reference genomes see Methods, Figure 5, and Dataset S2.

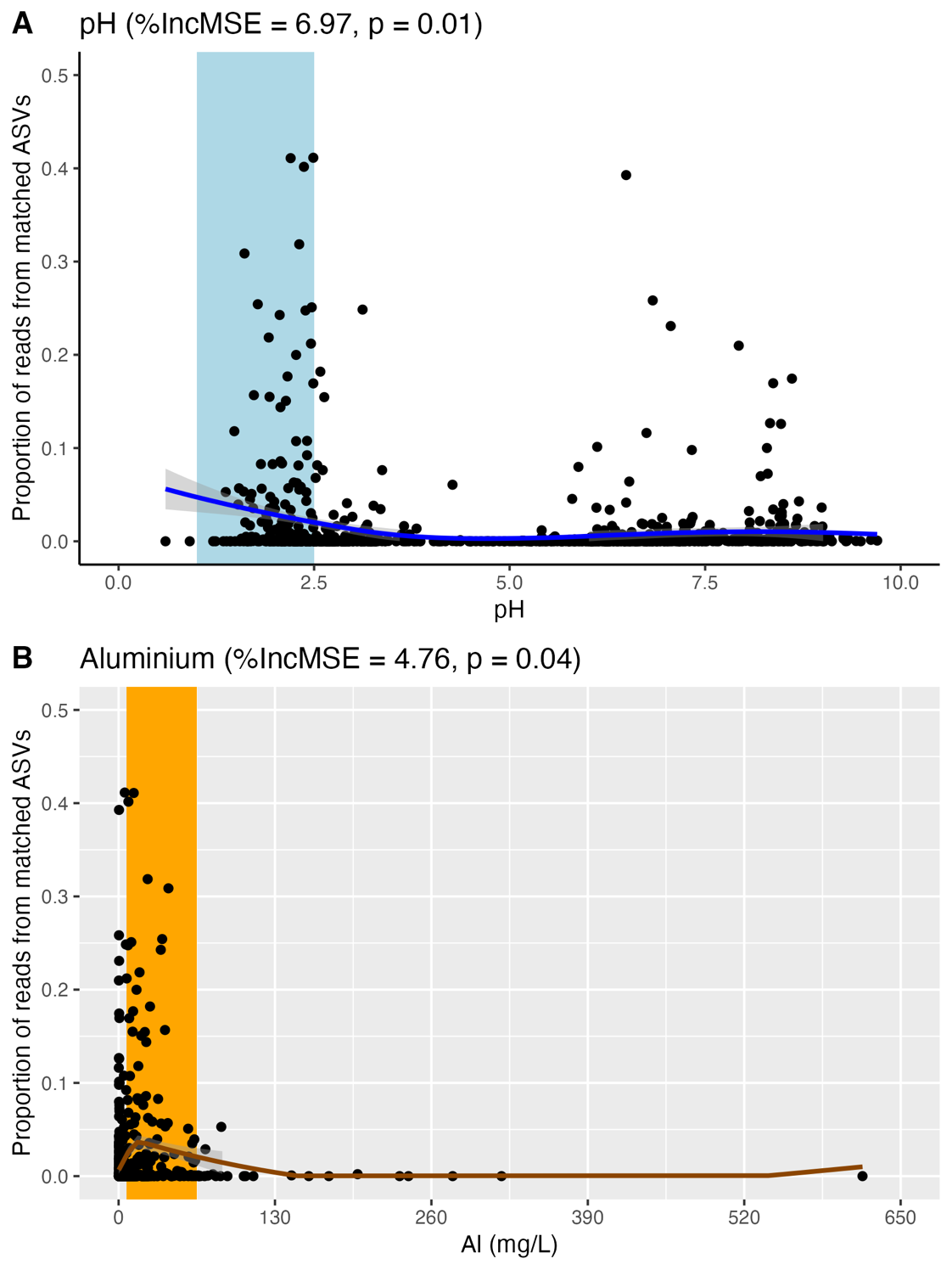

**Figure S8.** **A)** The results of the predictive modeling for pH, the variable identified as being most significant to determining where thermophilic plastic degrading bacteria may be found. Line of best fit for the generalized additive model (GAM) is displayed in blue. Shaded area indicates the range in which thermophilic plastic degrading organisms are predicted to be the most abundant. **B)** The results of the predictive modeling for Aluminum (Al), which was significant (p = 0.04) but did not increase the mean standard error more than 5%. Line of best fit for the generalized additive model (GAM) is displayed. Shaded area indicates the range in which thermophilic plastic degrading organisms are predicted to be the most abundant. GAM metrics can be found in Table S4

**Dataset S1. (Separate File, Dataset_S1 _Incubations.xlsx)**

This file contains the information about the gyroids used for the **A)** multi-substrate incubations and **B)** time series incubations. For both experiments, spring conditions (temperature, conductivity, pH) at start of incubation are included as well as mass of gyroid before incubation and mass of gyroid after incubation.

**Dataset S2. (Separate File, Dataset_S2 _ASVTable.xlsx)**

This file contains the ASV table of all bacterial and archaeal table sequences recovered through our cultivation-independent sequencing of the biofilms shaken from the gyroids from the multi-substrate incubation. The 16S rRNA gene sequence associated with each ASV is listed.

**Dataset S3. (Separate File, Dataset_S3 _Isolates.xlsx)**

This file contains the information about the 2 PHB degrading isolates. **A)** information about the samples each organism was isolated from and the culturing conditions used. **B)** A.	Information about the genomes recovered from the seven plastic degrading isolates. including: details about the quality and completeness of each genome and the 16S rRNA gene sequence and the most similar accessions in GTDB and BLAST. **C)** Table of all plastic degrading and synthesis genes identified in the genomes based on the PlasticDB database (1). **D)** PHB degradation genes identified in the genomes.

**Dataset S4. (Separate File, Dataset_S4 _Predictions.xlsx)**

This file contains the data used for the predictive modeling of thermophilic plastic degraders. This includes **A)** The re-processed ASV table from the 1000 springs dataset (2), including 16S rRNA gene sequence for all recovered sequences. B) the 16S rRNA gene sequences of all thermophilic plastic degraders identified in our search of PlasticDB (1) (see methods for more details). Metadata associated with each sample from the 1000 springs project can be found in Power et al. (2).

**SI References:**

1. Gambarini V, Pantos O, Kingsbury JM, Weaver L, Handley KM, Lear G. 2022. PlasticDB: a database of microorganisms and proteins linked to plastic biodegradation. *Database* 2022:baac008.
2. Power JF, Carere CR, Lee CK, Wakerley GLJ, Evans DW, Button M, White D, Climo MD, Hinze AM, Morgan XC, McDonald IR, Cary SC, Stott MB. 2018. Microbial biogeography of 925 geothermal springs in New Zealand. *Nat Commun* 9:2876.
